# Chemistry-based vectors map the chemical space of natural biomes from untargeted mass spectrometry data

**DOI:** 10.1101/2025.01.22.634253

**Authors:** Pilleriin Peets, Aristeidis Litos, Kai Dührkop, Daniel R. Garza, Justin J.J. van der Hooft, Sebastian Böcker, Bas E. Dutilh

## Abstract

Untargeted metabolomics can comprehensively map the chemical space of a biome, but is limited by low annotation rates (<10%). We used chemistry-based vectors, consisting of molecular fingerprints or chemical compound classes, predicted from mass spectrometry data, to characterize compounds and samples. These chemical characteristics vectors (CCVs) estimate the fraction of compounds with specific chemical properties in a sample. Unlike the aligned MS1 data with intensity information, CCVs incorporate actual chemical properties of compounds, offering deeper insights into sample comparisons. Thus, we identified key compound classes differentiating biomes, such as ethers which are enriched in environmental biomes, while steroids enriched in animal host-related biomes. In biomes with greater variability, CCVs revealed key clustering compound classes, such as organonitrogen compounds in animal distal gut and lipids in animal secretions. CCVs thus enhance the interpretation of untargeted metabolomic data, providing a quantifiable and generalizable understanding of the chemical space of natural biomes.

## Main

Environmental systems contain a multitude of players including microbes^1,2^, viruses^3,4^, their encoded proteins, and metabolites^5,6^ that together need comprehensive multi-omics approaches to chart and understand^7^. Microbes are involved in both producing and consuming metabolites. While some metabolic traits are encoded on the genome, a huge part remains unknown and represents chemical “dark matter”^8,9^. Besides metabolites, natural environments can contain other compounds, like abiotic or synthetic chemicals, which all combined create the chemical space. In environmental samples, the chemical space can be investigated by using nuclear magnetic resonance (NMR)^10,11^ or mass spectrometry (MS) coupled with liquid chromatography (LC)^12^ or gas-chromatography^13,14^. While NMR offers advantages in quantifying and identifying compounds, it falls short in sensitivity compared to MS, which emerged as the leading experimental platform in metabolomics^11^.

To chemically characterize samples, targeted^15^ and untargeted^16^ approaches are employed in LC-MS^12,17^. For precise identification and quantification of compounds, the targeted analysis uses previous knowledge about instrumental characteristics, such as retention time and fragmentation pattern, as well as chemical standards which may be expensive or difficult to obtain^17–19^. This approach can thus measure a limited number of dozens to hundreds^15^ of targeted compounds. For a more comprehensive overview of a sample, untargeted analysis using LC combined with high-resolution mass spectrometry (HRMS) is preferred, which can yield thousands of peaks for a single sample^20^. The acquisition of MS/MS fragmentation spectra is triggered in data-dependent acquisition for MS1 peaks (precursor peaks) with the highest intensity per time-point^21^. These fragmentation spectra aid in structure annotation through library matching with MS/MS spectra of known structures, or through networking analyses^22,23^.

The chemical composition of environmental and biological samples is highly complex and varies across biomes, making direct comparisons challenging. Current workflows often require the annotation of all detected compounds for meaningful comparisons, but this remains a bottleneck due to incomplete spectral libraries and the complexity of mass spectrometry (MS) fragmentation data. Platforms like *Inventa*^24^ have been developed for natural product discovery, aiming to identify plant extracts with higher potential for novel compounds. Similarly, tools like ReDU^25^, available within the Global Natural Products Social (GNPS) molecular networking platform, facilitate comparisons of chemical profiles with publicly available datasets. However, these approaches still emphasize comparisons based on annotated features, leaving a gap for robust sample comparisons based on broader chemical characterization.

The annotation of LC-MS peaks in metabolomics analysis can be challenging due to the abundance of peaks, incomplete spectral libraries, and the complexity of MS fragmentation data^20,26^. To determine compound structure, highly accurate mass-to-charge ratio (*m/z*) detection with information about the fragmentation pattern is essential. The GNPS molecular networking platform^22^ enables tentative annotation of up to 13% of LC-MS peaks^27^. Spectral libraries sometimes allow <5% of detected peaks to be structurally annotated, and even fewer to be identified via comparison to an in-house library^28,29^. Hence, the need arises for approaches prioritizing characterization over final structural identification, allowing for more insightful comparisons of samples even in the absence of complete annotations.

One characterization approach is to predict molecular fingerprints (MFP) and compound classes (CC)^30^, as done by using the SIRIUS+CSI:FingerID^31,32^ and CANOPUS^33^ software tools. Each bit of information in a MFP signifies the presence (1) or absence (0) of a specific substructure, including chemical elements, bonds, or functional groups. These characteristics allow for a nuanced understanding of the chemical composition of a complex sample without the necessity for exhaustive and often elusive final identification^31,32,34^. CSI:FingerID employs deep kernel learning to predict MFPs from MS/MS fragmentation data^31,35,36^. Instead of a binary fingerprint, CSI:FingerID predicts the probability that each substructure occurs in a compound. Besides adding probabilities to the MFP, SIRIUS also offers structural annotations^35^ and predicts CC probabilities^30,33,37^. Because MFP and CC reflect different structural aspects of the chemicals in a sample, using this information enables a more comprehensive characterization of the chemical space from untargeted data.

The Earth Microbiome Project (EMP) is a collaborative project analyzing microbial life from different biomes^1^. In their work with metabolomic datasets^9^, LC-MS peaks were assigned to one of eight categories (alkaloids, amino acids and peptides, carbohydrates, fatty acids, polyketides, shikimates and phenylpropanoids, terpenoids, or not annotated), and biomes were compared based on the relative abundance of these classes. Notably, biomes could be better differentiated when LC-MS intensity information was used, than when using presence/absence information alone. However, the inherent complexity of chemicals often means that assigning compounds to an individual category may not provide an accurate characterization. For example, assigning a compound to the alkaloids does not give information about other functional groups, such as the presence of sulfur in thiazole, or about the different number of rings in acridines and quinolines. All of this information is crucial for understanding the reactivity and biological activity of biochemical compounds.

In this study, we characterized the chemical space of samples at high resolution by employing molecular fingerprints and compound classes, enabling us to utilize LC-MS peaks that lacked definitive structural annotation. We developed a novel approach to compare samples efficiently by averaging the characteristics within each sample, creating a chemical characteristics vector (CCV) that describes the ratio of compounds with specific chemical characteristics. By comparing 572 EMP^9^ samples with the developed approach, we were able to distinguish samples from different biomes and identify biome-specific chemical features. We argue that CCVs provide a new way of understanding the chemical composition of natural samples.

## Methods

### Publicly available data

To test our model, we utilized data from the Earth Microbiome Project (EMP), which offers untargeted metabolomics datasets from a broad representation of biomes. Specifics about the samples and LC-HRMS methods are described in the EMP article^9^. We focused on 572 samples from eleven distinct biomes: animal corpus, animal distal gut, animal proximal gut, animal secretion, fungus corpus, plant detritus, plant surface, sediment, soil, subsurface, and water. Samples belonging to mixed biomes were excluded. For the biome annotation, we used the EMP Ontology (EMPO) level 4. To facilitate our analysis, we utilized the MS1 feature information file “1907_EMPv2_INN_GNPS_quant.csv” (downloaded project MSV000083475 from https://massive.ucsd.edu/), which includes necessary information about aligned LC-MS peaks and their intensities (peaks areas) in the corresponding samples.^9^

Absolute intensities in LC-MS do not directly correspond to compound concentrations due to numerous reasons, including differences in ionization efficiency between compounds^38,39^. Therefore, only differences in the ratio of LC-MS peak intensities across samples can provide meaningful information. In the comparative analysis of samples using MS1-based chromatographic peaks, we adopted two MS1 approaches for handling LC-MS peaks. Firstly, LC-MS peak intensities were discretized into binary presence/absence values. Alternatively, a logarithmic transformation (base 10) was applied to the intensities to standardize several magnitude differences in peak intensity. To ensure a fair comparison with the rest of the methods using MS/MS data, only peaks falling within the precursor *m/z* range of 100-900 were considered. This restriction resulted in a curated dataset consisting of 56,674 peaks. Uniform features with only zeros or other identical values were filtered out.

### Chemical characteristics calculations and preprocessing for sample comparison

For the CCV approach either multiple sample aligned data or individual LC-MS peak table is needed (Figure 1a). Preprocessing for feature extraction and alignment was done with SIRIUS 4.14 version. For LC-MS peaks with MS/MS data, we calculated molecular fingerprints (MFP) with CSI:FingerID^31,32,36^ an compound classes (CC) with CANOPUS^30,33^ inside SIRIUS software. Allowed elements for formula prediction were CHNOPS as well as Cl, Br, B, and Se based on the isotope pattern. Allowed mass deviation was 10 ppm or 0.002 Da. All probabilistic MFPs and CCs resulting from SIRIUS calculations were transformed to binary values using a threshold of 0.5 to indicate the presence or absence of specific chemical characteristics in the compound structures (Figure 1b). MFP and CC numbering is based on absoluteIndex from SIRIUS. MFP and CC files with descriptions for the workflow are available from Zenodo (see Data and code availability section).

**Figure 1.**
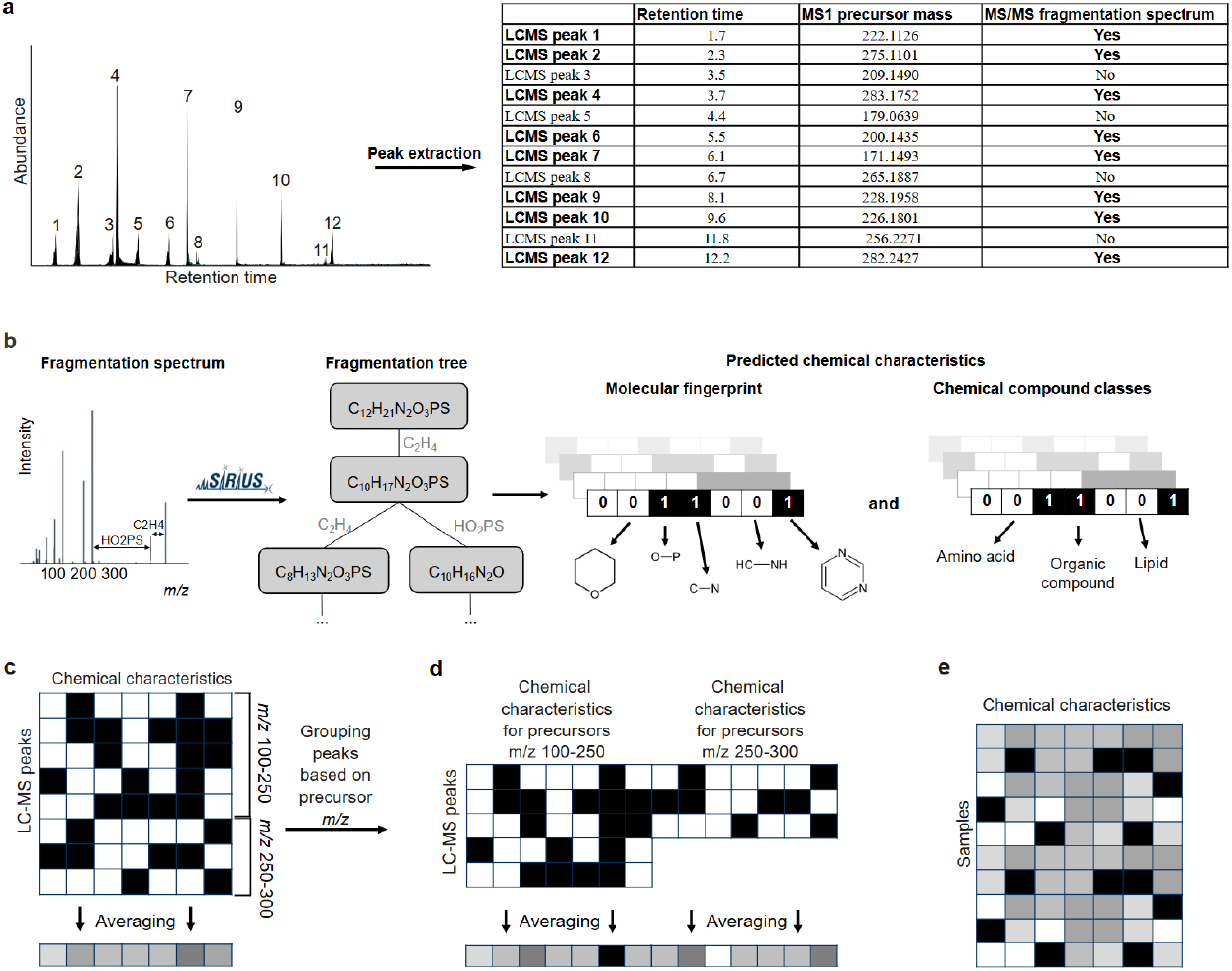
Workflow for comparing the chemical space of samples based on LC-MS data. (a) Peak extraction and filtering from raw LC-MS data, with key characteristics obtained in a feature quantification table, *i.e*., retention time, precursor mass, and whether or not MS/MS spectrum of this LC-MS peak was obtained. (b) SIRIUS to predict molecular fingerprints and compound class vectors for each LC-MS peaks with fragmentation spectrum. All probabilistic values from SIRIUS were transformed to binary with a threshold of 0.5. (c) Next, we calculated the fraction of compounds in a sample having a given substructure by averaging each column across all chemical features or (d) after grouping LC-MS peaks based on their precursor mass (100-250, 250-300, 300-350, 350-400, 400-450, 450-550, and 550-900 *m/z*) where the seven averaged vectors were concatenated. (e) The resulting matrix of averaged chemical characteristics vectors, generated by averaging all the chemical characteristics vectors for a sample, enables comparisons between samples.

The number of LC-MS peaks representing the chemical space varies depending on sample complexity (number of peaks per sample in Figure 2a), making direct between-sample comparisons challenging. Therefore, a standardization step was required. Here we averaged the chemical characteristics (Figure 1b) of peaks across all profiled MS/MS spectra to assess their overall prevalence in the sample. This results in a chemical characteristics vector (CCV) describing the fraction of compounds in a sample having a feature or substructure (Figure 1c). Besides averaging MFP and CC over all LC-MS peaks in the entire sample (Figure 1c), we also grouped the peaks based on their precursor *m/z* values before averaging (Figure 1d). For this, we established seven *m/z* groups to ensure an even distribution of peaks across samples: 100-250, 250-300, 300-350, 350-400, 400-450, 450-550, and 550-900 *m/z*.

**Figure 2.**
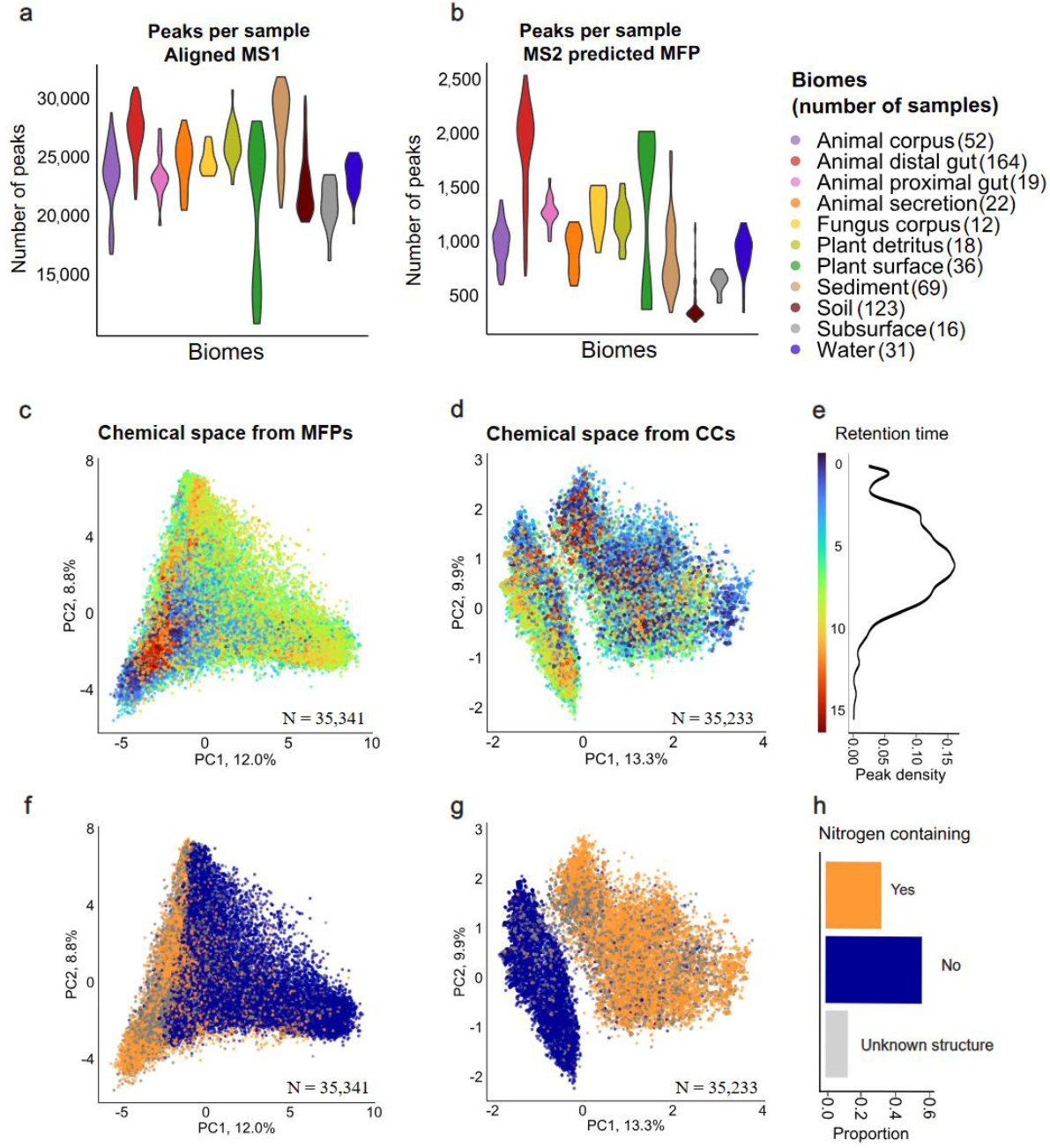
Evaluation of the LC-MS detected chemical space. Sample size distributions based on detected (a) LC-MS peaks with MS1 information and (b) MS/MS-based MFP information. (c) PCA plot based on MFP vectors for 35,341 LC-MS peaks in the EMP dataset colored by retention time (RT) and (d) based on SIRIUS CC vectors for 35,236 LC-MS peaks. Points are layered to highlight the less-represented RT groups, with values rounded to whole numbers. (e) Color gradient for plots c and d with compound distribution based on retention time. (f) Same PCA plot as on c, but colored by annotated nitrogen in the structure, and (g) same PCA as d, but colored by annotated nitrogen in the structure. (h) Color legend for plots f and g with information about the proportion of compounds with nitrogen presence in the predicted chemical structure.

### Modelling and statistical analysis

Downstream modeling and statistical analysis were conducted using R version 4.2.2 and Python version 3.10 for supervised machine learning (all code is available in GitHub). To identify important chemical characteristics from MFPs or CCs that contribute to sample classification on biomes, we employed a Recursive Feature Elimination (RFE)^44^ with the *rfe* function from the *caret* package in R. Specifically, we selected features in two separate sets, by first using elbow method^40^ selecting the optimal number of features (N) from validation sets (Table 1). As a first set, we obtained every feature that occurs at least once in the 10-fold cross-validation, which we refer to as rfe1. For rfe2, we only selected features that occur consistently in every fold in the cross-validation (Table 1).

**Table 1.** Final datasets for the analysis. Features with identical values for all samples were eliminated. RFE was applied to identify important features for biome discrimination. The rfe1 dataset consists of features that were selected in at least one of the 10 folds of cross-validation, while the rfe2 set contains features selected in all. A number of features (column three) were selected based on the elbow method^40^ on the feature importance.

| Feature | LC-HRMS peak grouping | N features for rfe | All features, eliminated zeros/ones | All important features (rfe1) | Consistently important features (rfe2) |
| --- | --- | --- | --- | --- | --- |
| LC-MS peak | Presence/absence | 1,000 | 56,646 | 4,407 | 42 |
|  | log10(Intensity) | 2,000 | 56,672 | 9,192 | 94 |
| Fingerprint | Averaged | 150 | 5,310 | 415 | 23 |
|  | Mass group averaged | 500 | 27,689 | 1,887 | 33 |
| Compound class | Averaged | 120 | 1,327 | 224 | 52 |
|  | Mass group averaged | 300 | 5,002 | 680 | 79 |

To assess to what extent our chemical representations captured the metabolomic differences between natural biomes, we utilized Principal Component Analysis (PCA) and Uniform Manifold Approximation and Projection (UMAP) for visualization. These dimensionality reduction techniques aid in visualizing the distribution and relationships among samples. Biome separation based on the aforementioned methods was validated by clustering metrics, such as the Calinski-Harabasz Index (CH), Dunn Index, and Silhouette index (Sil) (cluster.stats() function of the fpc package). To calculate these metrics, we used the biome annotation as cluster assignments and all the aforementioned methods as clustering input. CH assesses the compactness and separation of clusters, the Dunn Index measures the ratio of inter-cluster to intra-cluster mean distances, and the Silhouette index evaluates the cohesion and separation of clusters. These metrics collectively provide insights into the efficacy and performance of the methods employed for biome separation of the samples. For distance calculations, we used the Euclidean method in dist() function in R.

Moreover, we compared different chemical characteristic vector methods using supervised machine-learning techniques. We evaluated five binary classifiers: (i) Random Forest^41^ with 100 trees (number of estimators) and the Gini criterion (ii) k-nearest neighbors with k=5 and uniform weights per class^42^ (iii) Naive Bayes without prior probabilities^43^ and two trivial classifiers including (iv) the random classifier with equal probability for all classes, and (v) a classifier that predicts the most frequent class. We performed a k-fold cross-validation, where hyperparameters are selected based on the performance of the classifiers on a subset of the training data, when trained with the rest of the training dataset. This technique is common in classification tasks to optimize all parameters before applying the models to the validation set.

## Results

We set out to assess and compare the chemical composition of samples from different biomes based on high-resolution mass spectrometry data. We used data from the Earth Microbiome Project (EMP), which analyzed over 500 samples from 11 biomes^9^. We employed various methods to make samples comparable utilizing MS1 precursor data with intensities and MS/MS fragmentation data for predicted chemical characteristics information. We used fragmentation data from LC-MS to calculate averaged chemical characteristics vectors (CCVs) to describe the chemical space in each sample with a comparable data format. CCV describes the fraction of compounds in the LC-MS/MS data that were predicted to contain each specific characteristic (Figure 1c-d). This format allows adding sample data from different measurements, because it does not need aligned LC-MS data. Besides our newly developed approach implementing averaged molecular fingerprints (MFPs) and compound classes (CCs), we also included the aligned LC-MS peak table data with corresponding intensities for comparison^9^. MS1 approach was used as a benchmark method with the original detected peaks and their intensities. We performed a thorough comparison of the implemented approaches and revealed compound classes associated with biomes.

### Detected chemical space by LC-MS analysis

To compare the different biomes, we first evaluated the chemical space that was covered by the LC-MS measurements. In LC-MS analysis, this is limited to compounds with sufficient ionization efficiency^38^, and polarity in the range determined by the chromatographic column^44^. Detection of compounds in samples like soil and sediment additionally depends on the solvents used for extraction^45,46^. Finally, not all MS1 chromatographic peaks include MS/MS data, which is key for further annotation and chemical characterization of compounds. Altogether, 56,672 chromatographic peaks and their intensities (peak areas) were used to describe the chemical space of the 572 EMP samples. For MS/MS data, we conducted separate data processing and alignment with the same raw LC-MS data, obtaining 35,341 molecular fingerprint (MFP) vectors (62.4%) and 35,233 compound class (CC) vectors (62.2%). CSI:FingerID structurally annotated 30,795 (54.3%) of those peaks, including 4,145 (7.3%) annotated with high confidence (confidence score for the structure above 0.5)^36^. Notably, MFPs allow us to use all 35,341 peaks with MS/MS data, regardless of confidence scores for structural annotations, including those that did not receive a confident structural annotation. Each molecular fingerprint vector consists of 5,899 bits, and each compound class vector of 2,723 bits, each describing structural aspects of the compound.

Figures 2a and 2b show the number of processed chromatographic peaks, and MFPs predicted from acquired MS/MS data, respectively. In Figure 2b, a peak is only counted if MS/MS data was acquired from this specific sample. Although for this work aligned data was obtained for reducing the computational load, picking only peaks with MS/MS data per sample is imitating an alignment-free condition. With this alignment-free approach, less than 10% of the chromatographic peaks with MS1 detection have a corresponding MFP prediction.

To assess which components of the chemical space are captured by the MFP and CC vectors, we first projected all features from all the samples into two dimensions (Figure 2c,d,f,g). In Figure 2c, peaks with mid-range retention times spread across the plot, while peaks with low (RT < 4) or high (RT > 9) were clustered in the lower left corner. Compounds had normal distribution based on the retention time (Figure 2e). Retention time (RT) positively correlates with predicted octanol/water partition coefficient log*P*^47,48^, indicating that hydrophilic compounds have lower retention times (Supplementary Figure S1), which is in line with expectations of using C18-column based chromatography. As SIRIUS does not use retention time information when assigning structural annotation, this positive correlation indicates the correct annotation trend. For CC two distinct groups formed, with compounds containing nitrogen grouping separate from other compounds (Figure 2g). For MFP, some separation based on nitrogen presence was also seen (Figure 2f), but not so distinctly as for CC approach. MFP and CC vectors reveal that the chemical space is structured around hydrophobicity and the presence of nitrogen. On Supplementary Figures S2 and S3 compounds are additionally analyzed based on precursor mass-to-charge ratio. While the analysis of chemical characteristics (MFP and CC vectors) for individual compounds gives us an overview of the detected chemical space, to effectively compare the chemical space of samples, the amount of information needs to be reduced. We therefore proposed an averaging approach as introduced and discussed in the next sections.

### CCVs capture biome-specific chemical signatures with reduced features

To illustrate the relations between the samples and evaluate chemical characteristics vector (CCV) and MS1 approaches on their ability to separate samples from different biomes, we used two-dimensional principal component analysis (PCA) and Uniform Manifold Approximation and Projection (UMAP). While PC1 described around 20% of the variance in the MS1 data, over 50% of the variance is captured in those of the CCVs (Figure 3a-c). All three approaches grouped samples from similar biomes. For example, environmental biomes like soil, sediment, subsurface, and water grouped together with all approaches. However, we also observed a lot of mixing. Focusing on the UMAPs (Figure 3d-f), animal distal gut samples were generally divided into two major groups, with a few remote samples remaining. The soil samples formed two distinct groups in the MS1 plot (Figure 3d), while these samples were found closer together in the MFP-CCV plot (Figure 3e). Thus, to some extent, both MS1 and CCV approaches could distinguish biomes using chemical information obtained from the LC-MS measurements. PCA and UMAP figures for all other datasets are in Supplementary Table S1-S3.

**Figure 3.**
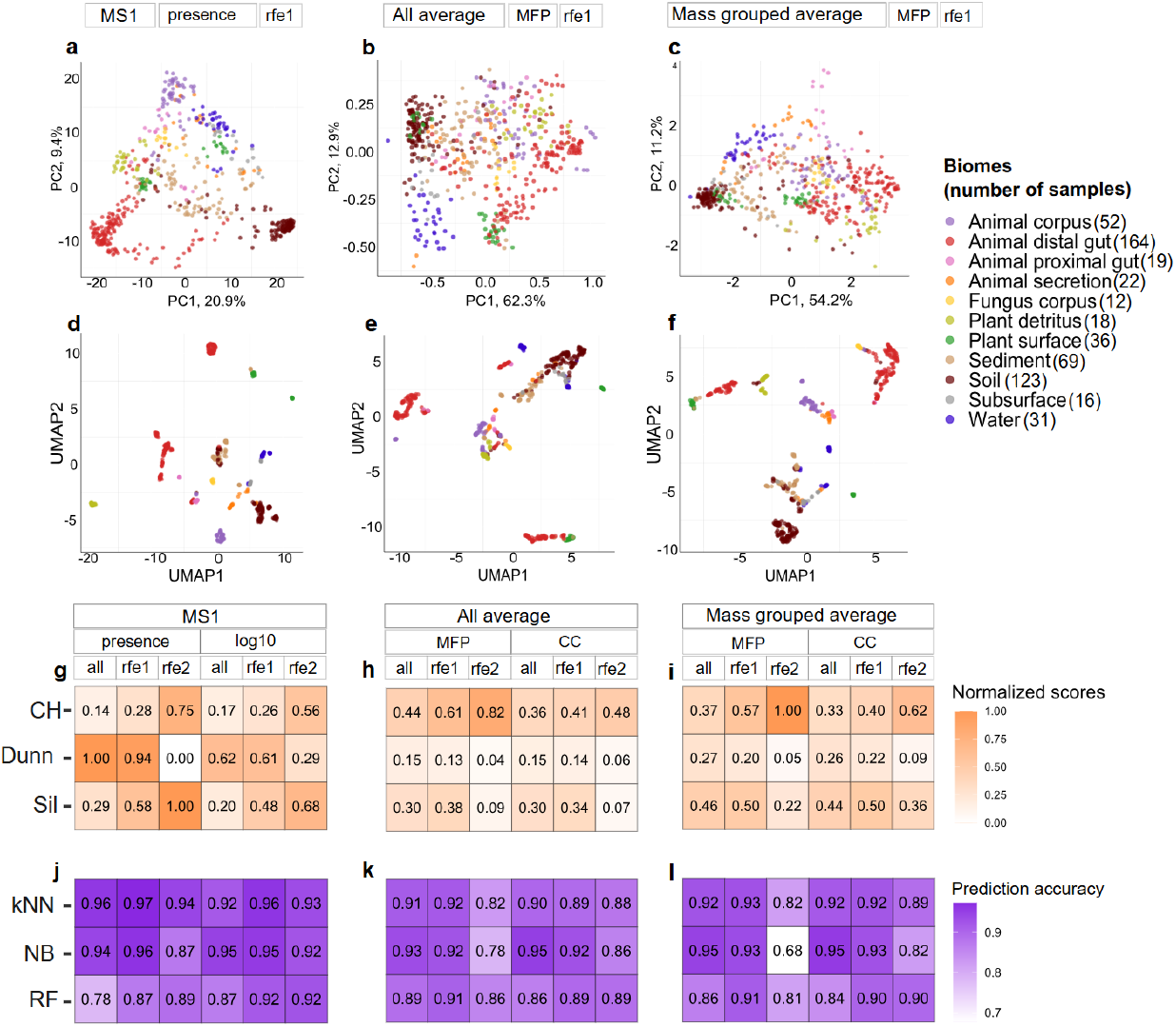
Separation of samples by biome according to various methods. (a) Each sample is characterized using LC-MS presence/absence data as input with selected important features (rfe1). (b) Selected sets of averaged fingerprints and (b) mass-grouped averaged fingerprints are used to describe each sample. (d-f) Corresponding UMAPs are shown for the same datasets as in a-c. Colors in (a-f) correspond to the sample biomes. (g-i) A comparison of 18 tested methods using clustering efficiency. In rows clustering metrics (normalized between 0 and 1), including Calinski-Harabasz (CH), Dunn, and Silhouette index (Sil). (j-l) The performance on biome prediction of the classifiers, k-Nearest Neighbors (kNN), Naive Bayes (NB), and Random Forest (RF). Methods are grouped based on the feature origin (MS1 peaks or chemical characteristics from MS/MS) and subsets selected by recursive feature elimination (rfe1: features that occurs at least once in the 10-fold cross-validation, rfe2: features that occur consistently in every fold in the cross-validation). PCA and UMAP graphs for the rest of the described methods are in Supplementary Table S1-S3 and the number of features for each approach in Table 1.

To quantify the quality of grouping of samples from different biomes (Figure 3g-i), we selected the Calinski-Harabasz (CH), Dunn, and Silhouette (Sil) indices to capture the compactness, separation, and cohesion of the biomes, respectively. The CH index quantifies the clustering performance in respect to the cluster centers, the Dunn index measures the separation between clusters, while Sil quantifies how close samples are within a cluster to the overall picture. Overall, MS1 approaches (Figure 3g) performed better than CCV (Figure 3h-i) in clustering the samples on their biome of origin. Interestingly, binary MS1 presence/absence data showed improved separation of the eleven biomes compared to logarithmically transformed values, suggesting that binary information enhanced the differences between clusters and samples. However, the different metrics varied substantially between the methods and did not provide uniform results. The Dunn index showed much higher values for MS^1^ than CCV approaches, whereas the CH index showed higher compactness values for the CCVs. This observation also holds when comparing the filtered datasets with recursive feature elimination (RFE)

The MFP and CC approaches, both derived from the MS data with SIRIUS predictions, resulted in fewer features than the MS1 approaches. Generally, more features correspond to higher clustering values, but MPF and CC approaches did not demonstrate a decrease in the clustering metrics. MFP outperformed CC, suggesting that they better captured biome-specific chemistry. Besides MFP having more features than CC, it also can describe compounds in more complex ways, compared to assigning compounds to defined compound classes. For example, molecular fingerprints can describe a very specific structural part, e.g., “does the compound have more than five aromatic rings?” or “does the compound have an aromatic ring with bromine group?”. Grouping the compounds into seven mass categories before averaging further improved performance (Figure 3i), showing the importance of this added resolution for chemically distinguishing biomes.

Feature selection had a big impact on the performance. The rfe1 dataset, where all features present in the 10-fold cross-validation were chosen, improved clustering, but the performance decreased with the rfe2 (dataset with constantly important features in 10-fold cross-validation), suggesting that overly stringent feature selection eliminated critical information about the chemical differences between the biomes.

Clustering indices did not unanimously agree on an optimal approach. The CH index improved with more restrictive datasets, rfe2 performing better than rfe1 and the full set of fingerprints in all cases, while the Dunn index showed a decrease in values, and the Silhouette index was less consistent. These results coincide with 2D plots, where we can see how samples from different biomes have many overlaps. These results highlight the value of using multiple clustering metrics to compare the methods in an unbiased manner.

### Supervised machine learning to compare the approaches

Next, we compared the MS1 and CCV methods in a supervised machine learning (ML) task to classify the samples by biome (e.g., animal distal gut, water, plant detritus). We utilized three non-linear algorithms, Random Forest (RF), k-nearest neighbors (kNN), and Naive Bayes (NB), as well as two trivial classifiers for reference, i.e., a random classifier where every biome is predicted with equal probability, and one that always predicts the most frequent biome in the training set (Figure 3j-l, Supplementary Table S4).

All ML methods had between 68-99% accuracy, compared to 9% and 29% for random with equal probability and most frequent classifiers, respectively (Supplementary Table S4). The probabilistic NB and kNN consistently performed better than RF. MS1 presence/absence dataset after feature selection including important features (rfe1 dataset) (Figure 3j) achieved the highest accuracy in kNN as well as NB classification. This was expected, as this data was binary without averaging, and therefore had less variability among the samples. Hence, kNN is able to find closer neighbors to the sample (looking at a presence/absence is easier than with values between 0 and 1). NB is probabilistic and assigns the probabilities of feature values to belong to a class, therefore it is easier to find more similar feature vectors for the validation samples. As a result, kNN was able to find closer neighbors for every validation sample, while NB was able to assign similar probabilities for the features of every validation sample.

Consistent with the clustering metrics, the 7-fold increased resolution of the mass-grouped data enhanced performance for all classification methods. MFP datasets had higher performance than CCs for RF and kNN, but not for the NB classifiers, as NB classifiers are more sensitive to outliers. Better results for MFP can be explained by a higher number of features as well as more informative characterization compared to compound class annotation. Overall, the classification results highlight the importance of feature selection, as well as the mass grouping of MS/MS datasets, on the final outcome.

### Sample origin influences the grouping within a biome

Above, we evaluated the methods based on their ability to distinguish and predict biomes, operating under the premise that samples from different biomes should exhibit distinct differences in their chemical space. However, as noted previously for microbiome DNA sequencing datasets^44^, manually defined biome annotations can be ambiguous, i.e., one annotation refers to multiple different systems, or redundant, i.e., different annotations reflect similar systems. We observed that some of the biomes split into several different clusters in the PCA and UMAP plots (Figure 3a-f). Such cases could either represent redundant biome labels referring to different systems, but it is also possible that they reflect batch effects or biases from sampling or analytical procedures. To address such biases systematically, we assessed whether samples from the same biome, but with different origins and/or contributors were still grouped together.

For animal distal gut, proximal gut, and plant surface samples, clustering often reflected the contributing study (Figure 4a,b). Samples originating from feces of different captive hosts, including mammals, birds, and herptiles (reptiles and amphibians), clustered remotely from the other animal distal gut samples, which included feces from Chernobyl voles, livestock, and whales from different contributors (Figure 4a). LC-MS peak amounts in these two clusters varied similarly, indicating that the bias is rather in the chemistry of the compounds and not in peak amounts. We also observed two distinct groups for the plant surface samples, which were derived from two different contributors and two different plant species (Figure 4b). This analysis shows that samples do not always have distinctive chemical space based on the biome annotation.

**Figure 4.**
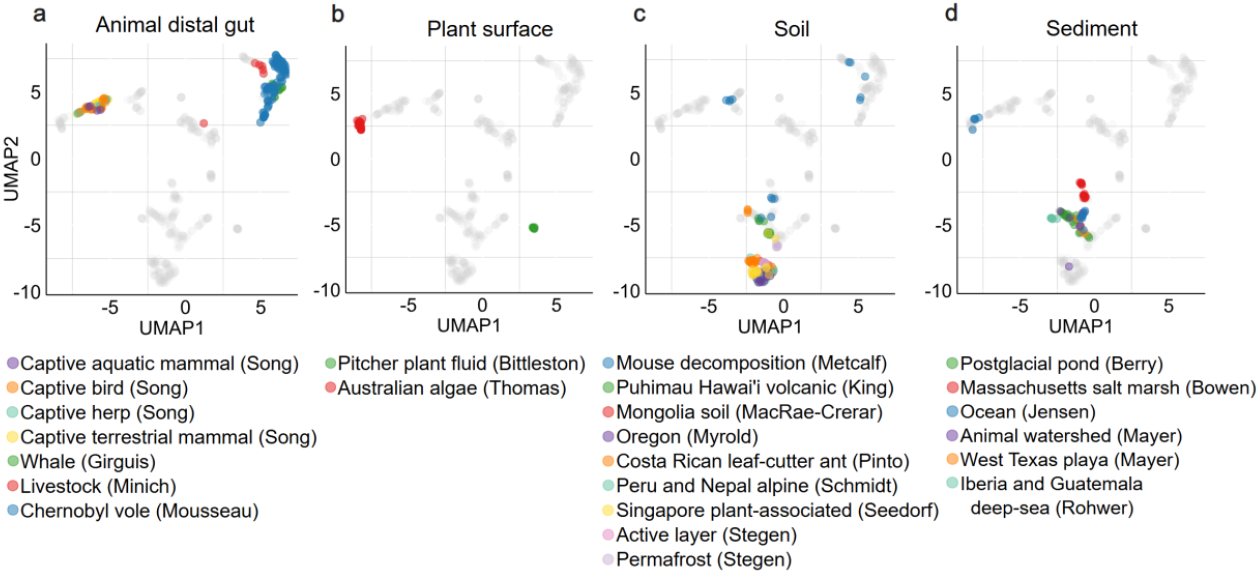
Two-dimensional projection of samples from different contributors and origins. UMAP plots were generated using rfe1-selected mass-grouped MFPs (as in Figure 3f). All 572 samples are shown in gray as the background for all plots. (a) Samples from different origins in the animal distal gut biome are highlighted in color. (b) Samples from different origins in the plant surface biome are highlighted in color. (c) Samples from different origins in the soil biome are highlighted in color. (d) Samples from different origins in the sediment biome are highlighted in color.

Soil and sediment samples consistently grouped together despite their diverse origin, except for two groups (Figure 4c,d). For soil, outliers represented mouse decomposition soil, while the other samples were regular soils from different origins. A similar trend can be seen for sediment samples (Figure 4d), where samples from five different contributors group together. The outliers from the Jensen group were samples from ocean sediments. There was a correlation for those outliers between LC-MS peak amounts and grouping, with outliers having more peaks per sample (e.g., for sediment samples with 1,800 peaks versus sediment samples <1,000 peaks).

All the animal secretion samples were generated by a single contributor, yet formed two separate groups in the UMAP plot (Figure 3d-f). Further metadata analysis revealed that these samples were collected from two different coral species from different locations (Panama, Western Australia). As we will see below, our CCV approach allows us to readily zoom into the compound classes that distinguish these two groups (Figure 6).

### Compound class ratios allow interpreting the chemical differences between biomes

To enhance our understanding of the chemical characteristics that distinguish different biomes, we examined the distribution of compound classes (CCs) among samples of different biomes. Unlike the aligned MS1 data, the chemical characteristics vector (CCV) approach incorporates actual chemical properties of compounds, offering deeper insights when comparing samples.

First, we looked into the compound classes themselves to assess their presence across different biomes. To identify specific chemical characteristics driving biome differentiation, we focused on the 52 most important compound classes (rfe2 dataset with consistently important features). The biome distributions and mean importance scores of all these 52 features are shown in Supplementary Figure S4. Most chemical characteristics were present in <10% of all detected LC-MS peaks (42/52 had an average <0.10) including specific classes like naphthols and furoic acid esters. Broader classes such as ethers and primary alcohols had higher averages. Here, we highlight seven key compound classes such as ethers, monosaccharide phosphates, glycerone phosphates, prenol lipids, steroids, bile acids, and amino acids (Figure 5). In upcoming sections, average values in CCV are multiplied by 100 to present the values as a % of LC-MS peaks per sample annotated with specific compound classes.

Ether bond, where oxygen is linked with two carbons, is common in secondary metabolites like carbohydrates, lipids, and terpenoids^49^. Ethers had the highest ratios among the 52 key features, with 7-44% of MS/MS features in each sample containing predicted ether bonds. This class was prominent in environmental biomes such as water (biome average of 33%), subsurface (30%), soil (25%), and sediment (25%). Biomes associated with animal hosts had ratio values below 22% (Figure 5a). Water, plant detritus, and fungal corpus biomes showed lower variability of sample-wise ratios than other biomes, suggesting better coincidence within the biome. Two distinct groups were observed within animal distal gut and plant surface biomes, matching the previous results from Figure 4a-b.

Monosaccharide phosphates and glycerone phosphates are organic compounds that play significant roles in biological systems, particularly in carbon and phosphorus cycling. Those phosphate compounds are considered intermediates involved in various critical metabolic pathways like monosaccharide phosphate in carbohydrate metabolism^50^. Animal, fungi, and plant detritus associated biomes consistently had low or near-zero levels of these compounds (Figure 5b,c), while plant surface and environmental biomes showed higher values and higher variability. Notably, soil samples exhibited a particularly wide range in glycerone phosphate values compared to other biomes, covering a >16-fold range spanning from 0.08-1.33% ratios. Subsurface and water samples had higher ratios than animal-associated biomes, ranging from 1-4% for monosaccharide phosphates and 0.09-0.75% for glycerone phosphates. Interestingly, distributions of the monosaccharides were more similar to the ether class in Figure 5a than monosaccharide phosphates. Monosaccharides in environmental biomes still showed high ratios, but higher ratios compared to monosaccharide phosphates were also noted for animal host-related biomes.

**Figure 5.**
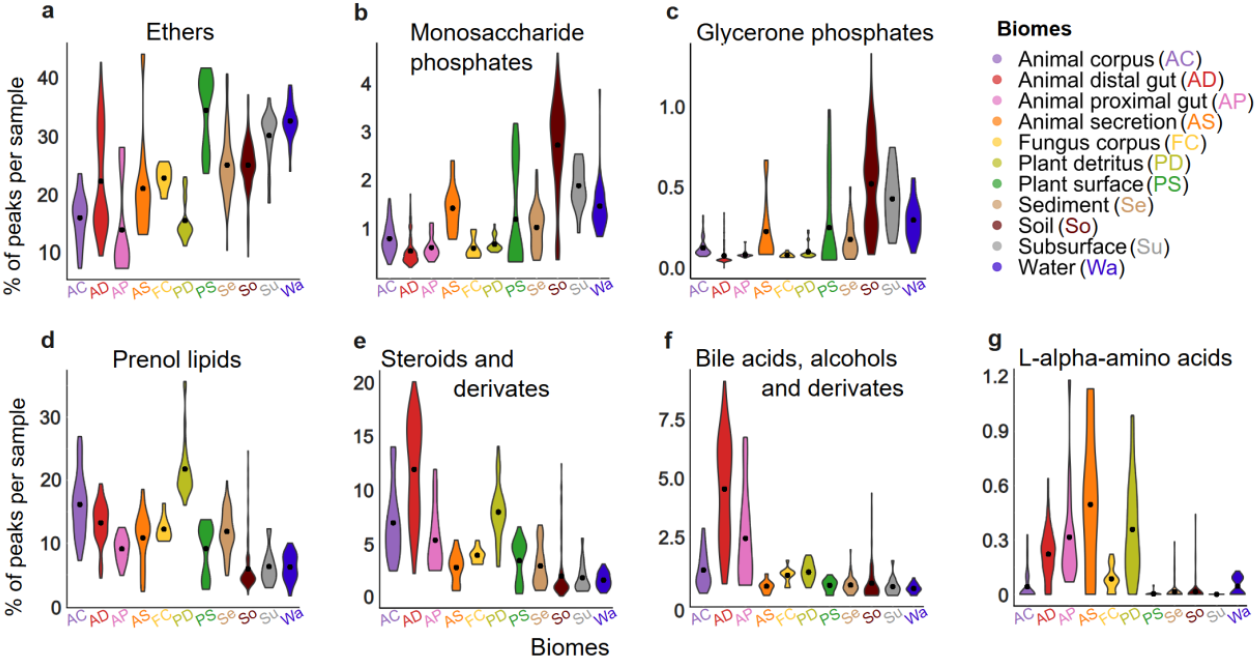
Characterization of biomes (x-axis) and samples using compound fractions (y-axis) in samples based on their compound class. Each violin plot on a graph shows the distribution of sample averages (average from chemical characteristics vector is multiplied with 100 to show the % instead of fraction) for the corresponding CANOPUS compound class predicted with SIRIUS+CSI:FingerID. Mean value for each biome is marked with black dot. The number of samples per biome are listed in Figure 2. (a) Distribution of compound class ratios for ethers in samples (b) Distribution of compound class ratios for monosaccharide phosphates in samples (c) Distribution of compound class ratios for glycerone phosphates in samples (d) Distribution of compound class ratios for prenol lipids in samples (e) Distribution of compound class ratios for steroids in samples (f) Distribution of compound class ratios for bile acids in samples (g) Distribution of compound class ratios for amino acids in samples.

In contrast to the ethers, monosaccharide phosphates, and glycerone phosphates described above, prenol lipids, steroids, bile acids, and amino acids were higher in animal-associated than in environmental samples (Figure 5d-g). While bile acids and amino acids had relatively low ratios, with the highest ratios reaching 9% (Figure 5f-g), prenol lipids and steroids with their derivatives reached ratios up to 36% and 20%, respectively (Figure 5d-e). Prenol lipids were the most frequent in plant detritus samples, ranging from 16-36%, while the values in other biomes stayed between 2-27%.

Steroids are derivatives of a cyclopentanophenanthrene 4-ring structure and are produced by humans, most other animals, and bacteria^51^. Natural steroids play crucial roles in various physiological processes, including stress response, immune response, carbohydrate metabolism, protein catabolism, regulation of blood electrolyte levels, and the control of inflammation and behavior^52^. For steroids and their derivates, the samples from animal corpus, distal and proximal gut, and plant detritus displayed wide variations and higher ratios compared to other biomes. For instance, in the animal distal gut, values ranged from 2-20% with a mean of 12%. In contrast, samples from water and subsurface biomes exhibited lower variability and lower values, with ranges between 0.3-3% and 0.4-5.4%, respectively (Figure 5e).

Bile acids and alcohols are a specific type of steroids found in mammals and other vertebrates, which are synthesized during cholesterol metabolism^53–55^. Indeed, high levels were observed in the animal corpus and gut biomes, while the rest of the biomes had near-zero values (Figure 5f). Still, even in the animal distal gut, only 0.4-9.2% (mean: 4.6 %) of compounds belonged to this compound class.

Alpha-amino acids are organic chemical compounds, containing both amino and carboxylic acid functional groups, and are used in nature as building blocks for proteins. Thus, a higher abundance of amino acids is expected in higher organisms such as animals (distal and proximal gut, secretion) and plant detritus, compared to environmental biomes (Figure 5g). Notably for the animal distal gut, the distribution is more dense, and for this compound class, we did not observe the two aforementioned separate groups. Animal corpus and fungus corpus, despite being higher organisms, had low amino acid content, and were more similar to environmental samples.

### Distribution of compound classes across mass groups

To assess the distribution of chemical characteristics in different compound sizes (molecular mass), we compared the seven selected compound classes using mass-grouped ratios (Supplementary Figure S5). For ethers, similar trends were observed across different mass groups. Higher ratios were noted for compounds of greater molecular mass, suggesting that heavier compounds had more ether bonds. Monosaccharide and glycerone phosphates were mostly present in higher molecular mass compounds, with glycerone phosphates only showing peaks at *m/z* > 550. In prenol lipids, particularly for plant detritus, higher average values were noted in higher mass groups. For alpha amino acids, different mass groups had very different values, with the highest ratios in mass group 450-550. The exceptions were animal proximal gut, distal gut, and plant detritus, where higher ratios were noticed in some lower mass groups as well. Using compound classes to characterize the clusters observed in the LC-MS data, highlights the key chemical traits that differentiate these biomes. Based only on the detected compounds with available MS/MS data, this approach offers valuable insights and enables meaningful comparisons of the chemical space of these samples in terms of interpretable compound categories.

### Compound classes explain differences within biomes: distal gut and coral samples

Animal distal gut samples showed wide variability in compound content, with 22/52 compound classes exhibiting greater spread than for other biomes. This was partly driven by the two distinct clusters seen in the 2D projections (Figures 3 and 4a). By describing the LC-MS data in terms of interpretable chemical characteristics, we could zoom into the chemical differences between these two groups. Group one, which included 54 samples contributed by the Song group, had many organoheterocyclic compounds (Figure 6a) and benzoyl derivatives (Figure 6b), while group two, which included 110 samples from three other contributors, had relatively many organonitrogen compounds (Figure 6c) and fatty amides (Figure 6d). For organonitrogen compounds (Figure 6c) we noticed a negative correlation in group two between the ratio of this compound class and the overall number of peaks per sample (Supplementary Figure S7). This indicates that abundance estimates could be artificially inflated if fewer other peaks were detected in a sample. When samples from groups one and two had similar numbers of peaks (>1,800 peaks, see Supplementary Figure S7), the compound class ratio values still showed significant differences between the two groups. This shows that separation into group one and group two was not related to the peak counts. The same observation was also confirmed for the other compound classes (Supplementary Figure S7).

**Figure 6.**
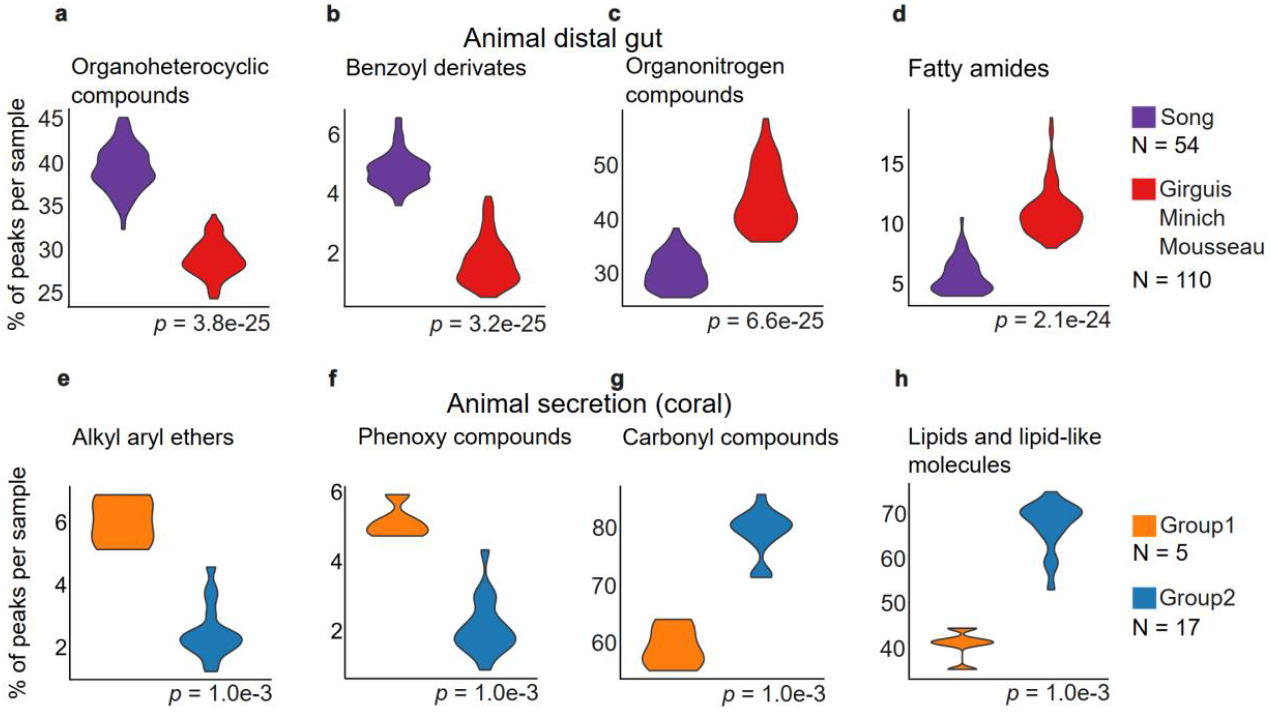
Within biome differences for sample clusters. (a-d) Four key compound classes to differentiate two distant animal distal gut sample clusters (Figure 4a). Song group has feces samples from captive birds, herps, and terrestrial and aquatic mammals.^1^ (e-h) Four key compound classes to differentiate two distant coral sample clusters (from Figure 3f, Supplementary Figure S7). Coral (*Pocillopora damicornis* and *Porites lobata*) samples originated from Panama and Western Australia.^1^ y-axis represents the % of compounds in each sample with specific compound class annotation. p-values are calculated using Wilcoxon rank-sum test.

We also observed two distinct groups in the animal secretion biome, which consisted of 22 coral samples from one contributor (Supplementary Figure S8). Here, metadata analysis revealed that these groups likely corresponded to two different coral species, sampled at two geographical locations^9^. Of the 58 key compound classes differentiating the two groups of secretion samples, 57% of CCs had higher ratios in group one than in group two (Supplementary Figure S9). Less than 10% of the peaks belonged to the 33 CCs associated with group one. This indicates that key classes for group one included specific annotations like alkyl aryl ethers (Figure 6e) and phenoxy compounds (Figure 6f). Group two was enriched for more generally annotated CCs like carbonyl compounds (Figure 6g) and lipids (Figure 6h).

### Limitations and future perspectives

The extent of the chemical space that can be detected by LC-MS depends on various factors. For instance, sample preparation, chromatographic separation, ionization efficiency, and MS/MS data acquisition all have their specific biases and limitation^46^. Thus, the compounds detected in a metabolomics survey cannot represent the full complexity of the chemical composition in nature. Nevertheless, untargeted metabolomics allows for a high-throughput, relatively unbiased comparison of samples and their chemical space.

In this work, we used peak alignment to reduce the calculation volume prior to SIRIUS analysis. However, we note that the CCV approach can be run independently of peak alignment, which is usually dependent on a consistent experimental setup. Thus, CCVs enable comparisons between samples that are measured later under different conditions. For untargeted LC-MS data, CCVs thus allow for cross-study comparison; however, such comparisons require careful analysis and comparison of the initial covered chemical space of the instrumental method.

One notable limitation is that the current approach only considers the presence or absence, and not the abundance of compounds in the sample. Due to the detection limit of the LC-MS machine and the noise cutoffs used in our analysis, only compounds higher than a minimum concentration will be detected and can be compared, and trace compounds are unlikely to be assessed. Consequently, the reported ratios of compound classes and structural parts are qualitative, reflecting the diversity of the detected compounds containing a given substructure, but not the absolute abundance or concentration of these compounds. In studies where samples are measured under consistent conditions and where internal standards are used for peak standardization, incorporating weights according to peak intensity can improve the results of chemical space comparisons by accounting for quantitative differences as well.

For future applications, we note that CCVs are not limited to comparing metabolomics samples derived from different (micro-) biomes, but can also be applied to identify chemical differences in environmental monitoring samples, time-series data, or comparisons of sick and healthy organisms. In multi-omics studies where both metabolomics and metagenomics measurements are conducted, chemical characteristic ratios can be correlated with microbes, as well as the presence or expression of specific genes or metabolic pathways, opening the potential for linking microbial traits to metabolomic features.

## Discussion

We developed a novel approach to characterize metabolomics data using a chemical characteristics vector (CCV) by leveraging SIRIUS-predicted molecular fingerprints and chemical compound classes. These chemical characteristics vectors enabled us to compare samples and biomes, as well as characterize their chemical space in terms of compound classes. To simulate alignment-free conditions, we utilized MS/MS data acquired from specific samples, describing approximately 10% of the detected data per sample using predicted chemical characteristics.

Using 572 samples assigned to 11 biomes from the Earth Microbiome Project (EMP), we demonstrated that averaged chemical characteristics could cluster biomes as effectively as benchmark LC-MS methods based on peak intensities. Grouping peaks by their precursor mass prior to averaging improved results for both molecular fingerprints and compound classes. In general, both machine learning-based predictions and clustering metrics improved. Since chemical characteristics vectors contain thousands of features across fewer than 600 samples, we showed that reducing the features with recursive feature elimination improved biome clustering. More stringent feature reduction decreased performance, as essential information about the chemical differences between the biomes was likely removed.

Beyond grouping the biome samples, we explored the compound classes shaping the chemical space in the different biomes. For example, in the animal gut, ethers represented <20% of the annotated LC-MS peaks, compared to >30% in water. Environmental samples such as water, soil, and sediment generally contained higher proportions of ethers, while steroids, bile acids, and amino acids were more dominant in animal-associated biomes.

While samples from the same biome were generally similar, several biomes exhibited significant variability. For example, the clustering of animal distal gut, plant surface, soil, and sediment samples depended heavily on their origin. Using our newly calculated averaged compound class values, we identified chemical differences between animal distal gut and coral samples, providing new insights into the chemical biology of these animals. Overall, our alignment-free LC-MS method facilitates interpretable chemical comparisons between samples, providing a versatile tool for exploring the metabolomic diversity of the natural world.

## Data and code availability

Data and the code are available on GitHub (https://github.com/MGXlab/MetabSpace) and Zenodo (https://doi.org/10.5281/zenodo.14506250).

## Supplementary Information

The supplementary PDF file “SupplementaryInformation1.pdf” contains additional UMAP and PCA figures, compound class plots for biomes, and a table with prediction model results.

## Acknowledgments

We thank the people in the Viral Ecology and Omics group at Jena University. We also thank Emma Schymanski and her group at the University of Luxembourg for their feedback and advice. Sample processing was performed by the Earth Microbiome Project (https://earthmicrobiome.org), and all the data and metadata have been made public through the EMP data portal (https://qiita.microbio.me/emp).

## Author Contributions

P.P., B.E.D., S.B., J.J.J.v.d.H., and D.G. conceived and designed the study. P.P. analyzed EMP data, and wrote the code for data analysis. A.L. wrote the code for the statistical comparison approach and ML models. K.D. conducted SIRIUS calculations. P.P., A.L., and B.E.D. wrote the paper with input from all authors. All authors read and approved the manuscript.

## References

1. Thompson, L. R. et al. A communal catalogue reveals Earth’s multiscale microbial diversity. Nature 551, 457–463 (2017).

2. Von Meijenfeldt, F. A. B., Hogeweg, P. & Dutilh, B. E. A social niche breadth score reveals niche range strategies of generalists and specialists. Nat. Ecol. Evol. 7, 768–781 (2023).

3. Paez-Espino, D. et al. Uncovering Earth’s virome. Nature 536, 425–430 (2016).

4. Gregory, A. C. et al. Marine DNA Viral Macro- and Microdiversity from Pole to Pole. Cell 177, 1109–1123.e14 (2019).

5. Loureiro, C., Medema, M. H., Van Der Oost, J. & Sipkema, D. Exploration and exploitation of the environment for novel specialized metabolites. Curr. Opin. Biotechnol. 50, 206–213 (2018).

6. Krautkramer, K. A., Fan, J. & Bäckhed, F. Gut microbial metabolites as multi-kingdom intermediates. Nat. Rev. Microbiol. 19, 77–94 (2021).

7. Eren, A. M. & Banfield, J. F. Modern microbiology: Embracing complexity through integration across scales. Cell 187, 5151–5170 (2024).

8. Navarro-Muñoz, J. C. et al. A computational framework to explore large-scale biosynthetic diversity. Nat. Chem. Biol. 16, 60–68 (2020).

9. Shaffer, J. P. et al. Standardized multi-omics of Earth’s microbiomes reveals microbial and metabolite diversity. Nat. Microbiol. 7, 2128–2150 (2022).

10. Emwas, A.-H. et al. NMR Spectroscopy for Metabolomics Research. Metabolites 9, 123 (2019).

11. Markley, J. L. et al. The future of NMR-based metabolomics. Curr. Opin. Biotechnol. 43, 34–40 (2017).

12. Gorrochategui, E., Jaumot, J., Lacorte, S. & Tauler, R. Data analysis strategies for targeted and untargeted LC-MS metabolomic studies: Overview and workflow. TrAC Trends Anal. Chem. 82, 425–442 (2016).

13. Zeki, Ö. C., Eylem, C. C., Reçber, T., Kır, S. & Nemutlu, E. Integration of GC–MS and LC–MS for untargeted metabolomics profiling. J. Pharm. Biomed. Anal. 190, 113509 (2020).

14. Beale, D. J. et al. Review of recent developments in GC–MS approaches to metabolomics-based research. Metabolomics 14, 152 (2018).

15. Zhou, J. & Yin, Y. Strategies for large-scale targeted metabolomics quantification by liquid chromatography-mass spectrometry. The Analyst 141, 6362–6373 (2016).

16. Klåvus, A. et al. “Notame”: Workflow for Non-Targeted LC–MS Metabolic Profiling. Metabolites 10, 135 (2020).

17. Russo, F., Ottosson, F., Van Der Hooft, J. J. J. & Ernst, M. Deep Learning Models for LC-MS Untargeted Metabolomics Data Analysis. in From Computational Logic to Computational Biology (eds. Cantone, D. & Pulvirenti, A.) vol. 14070 128–144 (Springer Nature Switzerland, Cham, 2024).

18. Büscher, J. M., Czernik, D., Ewald, J. C., Sauer, U. & Zamboni, N. Cross-Platform Comparison of Methods for Quantitative Metabolomics of Primary Metabolism. Anal. Chem. 81, 2135–2143 (2009).

19. Ribbenstedt, A., Ziarrusta, H. & Benskin, J. P. Development, characterization and comparisons of targeted and non-targeted metabolomics methods. PLOS ONE 13, e0207082 (2018).

20. Tian, Z., Liu, F., Li, D., Fernie, A. R. & Chen, W. Strategies for structure elucidation of small molecules based on LC–MS/MS data from complex biological samples. Comput. Struct. Biotechnol. J. 20, 5085–5097 (1011).

21. Guo, J. & Huan, T. Comparison of Full-Scan, Data-Dependent, and Data-Independent Acquisition Modes in Liquid Chromatography–Mass Spectrometry Based Untargeted Metabolomics. Anal. Chem. 92, 8072–8080 (2020).

22. Wang, M. et al. Sharing and community curation of mass spectrometry data with Global Natural Products Social Molecular Networking. Nat. Biotechnol. 34, 828–837 (2016).

23. Horai, H. et al. MassBank: a public repository for sharing mass spectral data for life sciences. J. Mass Spectrom. 45, 703–714 (2010).

24. Quiros-Guerrero, L.-M. et al. Inventa: A computational tool to discover structural novelty in natural extracts libraries. Front. Mol. Biosci. 9, 1028334 (2022).

25. Jarmusch, A. K. et al. ReDU: a framework to find and reanalyze public mass spectrometry data. Nat. Methods 17, 901–904 (2020).

26. Rathahao-Paris, E., Alves, S., Junot, C. & Tabet, J.-C. High resolution mass spectrometry for structural identification of metabolites in metabolomics. Metabolomics 12, 10 (2016).

27. Bittremieux, W., Wang, M. & Dorrestein, P. C. The critical role that spectral libraries play in capturing the metabolomics community knowledge. Metabolomics 18, 94 (2022).

28. Panagopoulos Abrahamsson, D. et al. A Comprehensive Non-targeted Analysis Study of the Prenatal Exposome. Environ. Sci. Technol. 55, 10542–10557 (2021).

29. Newton, S. R. et al. Suspect screening and non-targeted analysis of drinking water using point-of-use filters. Environ. Pollut. 234, 297–306 (2018).

30. Djoumbou Feunang, Y. et al. ClassyFire: automated chemical classification with a comprehensive, computable taxonomy. J. Cheminformatics 8, 61 (2016).

31. Dührkop, K. et al. SIRIUS 4: a rapid tool for turning tandem mass spectra into metabolite structure information. Nat. Methods 16, 299–302 (2019).

32. Heinonen, M., Shen, H., Zamboni, N. & Rousu, J. Metabolite identification and molecular fingerprint prediction through machine learning. Bioinformatics 28, 2333–2341 (2012).

33. Dührkop, K. et al. Systematic classification of unknown metabolites using high-resolution fragmentation mass spectra. Nat. Biotechnol. 39, 462–471 (2021).

34. O’Boyle, N. M. & Sayle, R. A. Comparing structural fingerprints using a literature-based similarity benchmark. J. Cheminformatics 8, 36 (2016).

35. Dührkop, K., Shen, H., Meusel, M., Rousu, J. & Böcker, S. Searching molecular structure databases with tandem mass spectra using CSI:FingerID. Proc. Natl. Acad. Sci. 112, 12580–12585 (2015).

36. Hoffmann, M. A. et al. High-confidence structural annotation of metabolites absent from spectral libraries. Nat. Biotechnol. 40, 411–421 (2022).

37. Kim, H. W. et al. NPClassifier: A Deep Neural Network-Based Structural Classification Tool for Natural Products. J. Nat. Prod. 84, 2795–2807 (2021).

38. Leito, I. et al. Towards the electrospray ionization mass spectrometry ionization efficiency scale of organic compounds. Rapid Commun. Mass Spectrom. 22, 379–384 (2008).

39. Tang, X., Bruce, J. E. & Hill, H. H. Characterizing Electrospray Ionization Using Atmospheric Pressure Ion Mobility Spectrometry. Anal. Chem. 78, 7751–7760 (2006).

40. Humaira, H. & Rasyidah, R. Determining The Appropiate Cluster Number Using Elbow Method for K-Means Algorithm. in Proceedings of the Proceedings of the 2nd Workshop on Multidisciplinary and Applications (WMA) 2018, 24-25 January 2018, Padang, Indonesia (EAI, Padang, Indonesia, 2020). doi:10.4108/eai.24-1-2018.2292388.

41. Breiman, L. Random Forest. Mach. Learn. 45, 5–32 (2001).

42. Cover, T. & Hart, P. Nearest neighbor pattern classification. IEEE Trans. Inf. Theory 13, 21–27 (1967).

43. Zhang, H. The Optimality of Naive Bayes. in The Optimality of Naive Bayes vol. (FLAIRS 2004) (The AAAI Press, Menlo Park, California, 2004).

44. Hemmer, S., Manier, S. K., Wagmann, L. & Meyer, M. R. Comparison of reversed-phase, hydrophilic interaction, and porous graphitic carbon chromatography columns for an untargeted toxicometabolomics study in pooled human liver microsomes, rat urine, and rat plasma. Metabolomics 20, 49 (2024).

45. Brown, R. W. et al. Soil metabolomics - current challenges and future perspectives. Soil Biol. Biochem. 193, 109382 (2024).

46. Black, G. et al. Exploring chemical space in non-targeted analysis: a proposed ChemSpace tool. Anal. Bioanal. Chem. 415, 35–44 (2023).

47. Leo, A., Hansch, C. & Elkins, D. Partition coefficients and their uses. Chem. Rev. 71, 525–616 (1971).

48. Leo, A. J. Calculating log Poct from structures. Chem. Rev. 93, 1281–1306 (1993).

49. Domínguez De María, P., Van Gemert, R. W., Straathof, A. J. J. & Hanefeld, U. Biosynthesis of ethers: Unusual or common natural events? Nat. Prod. Rep. 27, 370 (2010).

50. Bhagavan, N. V. Medical Biochemistry. (Harcourt/Academic Press, San Diego, 2002).

51. Ying, G.-G., Kookana, R. S. & Ru, Y.-J. Occurrence and fate of hormone steroids in the environment. Environ. Int. 28, 545–551 (2002).

52. Rasheed, A. & Quasim, Mohd. A Review of Natural Steroids and their Applications. Int. J. Pharm. Sci. Res. 4, 520–531 (2013).

53. Hofmann, A. F., Hagey, L. R. & Krasowski, M. D. Bile salts of vertebrates: structural variation and possible evolutionary significance. J. Lipid Res. 51, 226–246 (2010).

54. Quinn, R. A. et al. Global chemical effects of the microbiome include new bile-acid conjugations. Nature 579, 123–129 (2020).

55. Chiang, J. Y. L. Bile Acid Metabolism and Signaling. in Comprehensive Physiology (ed. Terjung, R.) 1191–1212 (Wiley, 2013). doi:10.1002/cphy.c120023.

