## Supplementary Information 1 for "Chemistry-based vectors map the chemical space of natural biomes from untargeted mass spectrometry data"

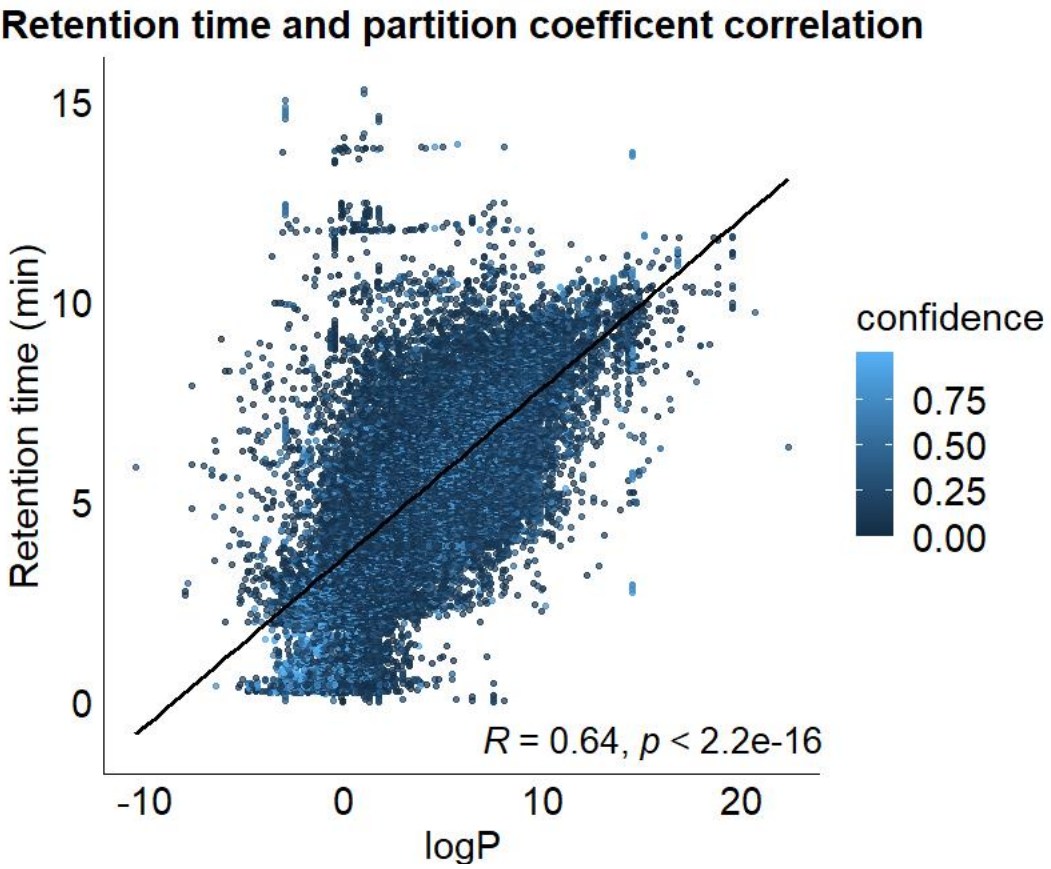


**Figure S1.** Correlation between measured retention time and octanol-water partition coefficient log*P* predicted from the rank1 structural annotation. Points are colored based on SIRIUS assigned confidence score for the rank1 structure. For this data analysis, reverse phase chromatography was used and thus positive correlation shows that the predicted compounds follow the expected correlation that more polar compounds with low logP value exit the column earlier, while compounds with high log*P* exit it later having high retention time. As this is the expected result, having a positive correlation confirms the correct prediction of structural annotations done by SIRIUS+CSI:FingerID software.

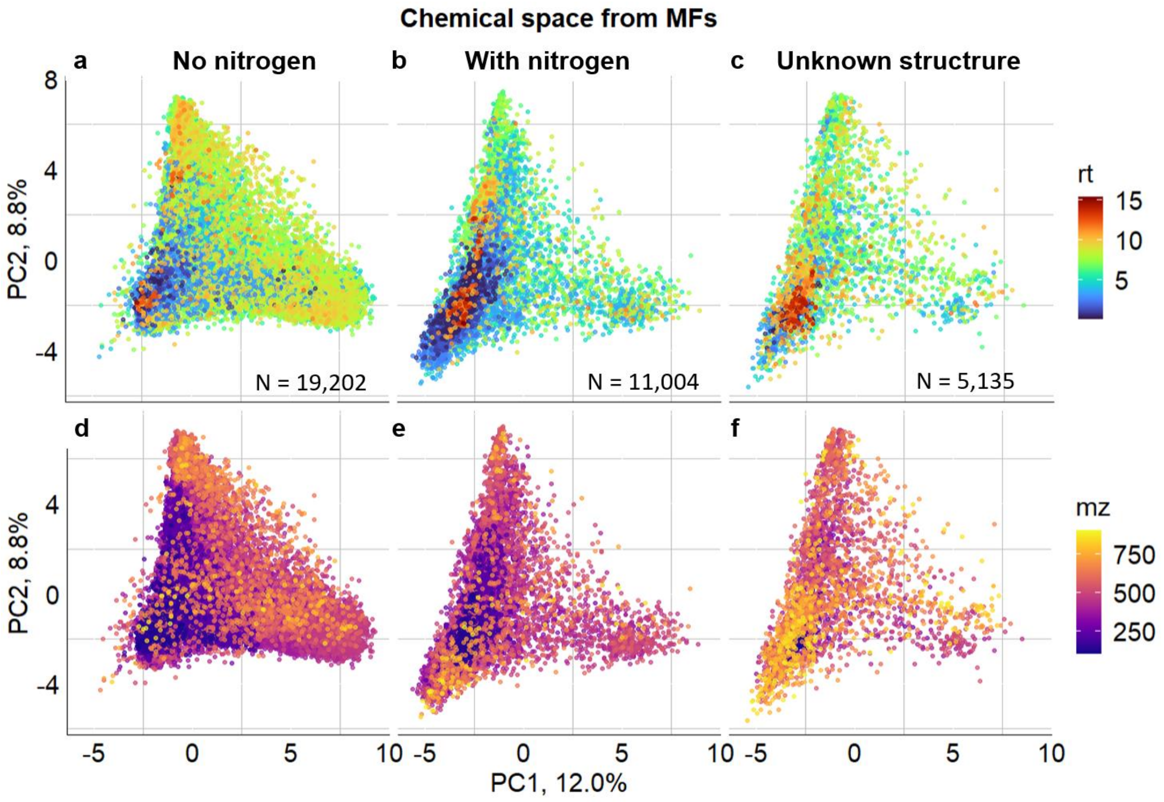


**Figure S2.** Detected chemical space projection using molecular fingerprints (MFPs). Here, the main text Figure 2c is divided into three to show the separation of points better. (a-c)The coloring of the points is based on experimental retention time (rt) and (d-f) based on the precursor mass-to-charge ratio (mz). Separation of points for the plot based on the (a,d) absence of nitrogen and (b,e) presence of nitrogen in the predicted structure. (c,f) LC-MS features without structural annotation.

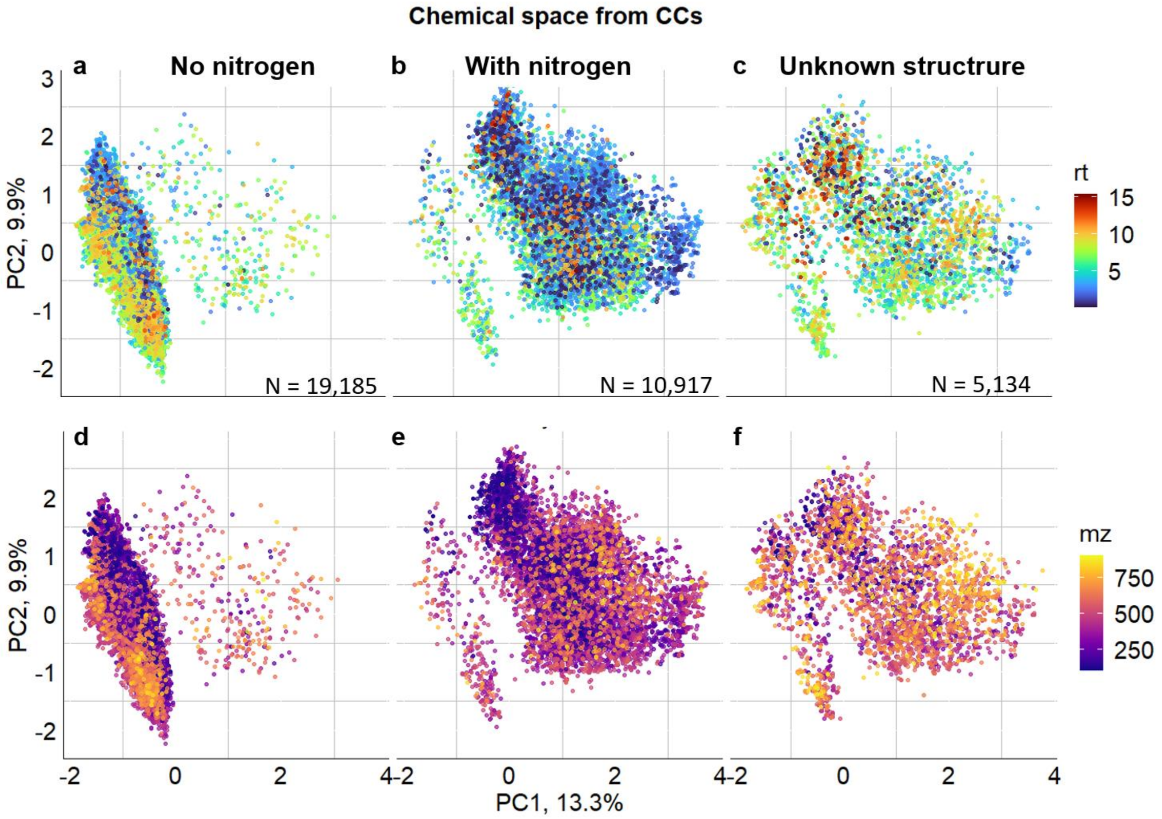


**Figure S3.** Detected chemical space projection using compound classes (CCs). Here the main text Figure 2d is divided into three to show the separation of points better. (a-c) The coloring of the points is based on experimental retention time (rt) and (d-f) based on the precursor mass-to-charge ratio (*mz*). Separation of points for the plot based on the (a,d) absence of nitrogen and (b,e) presence of nitrogen in the predicted structure. (c,f) LC-MS features without structural annotation.

**Table S1.** Principal component analysis (PCA), Uniform Manifold Approximation, and Projection (UMAP) for sample comparisons for MS1 datasets. Aligned LC-MS features with corresponding presence/absence or intensity information in log10 form were used. For both approaches, datasets were analyzed with all features (marked as all), selecting all important features from recursive feature elimination 10-fold cross-validation (rfe1) and selecting consistently important features present in all 10-fold cross-validation when using recursive feature elimination (rfe2).

| Method | PCA | UMAP |
|---|---|---|
| MS1<br>presence/absence<br>all | 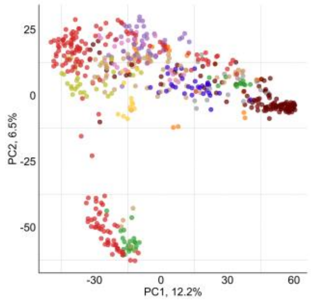 | 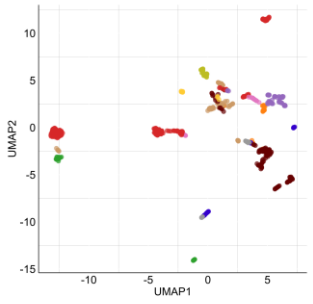 |
| MS1<br>presence/absence<br>rfe1 | 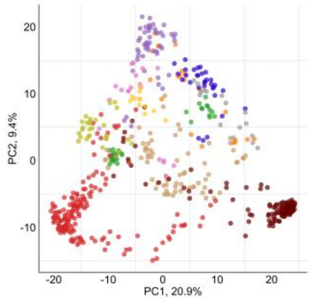 | 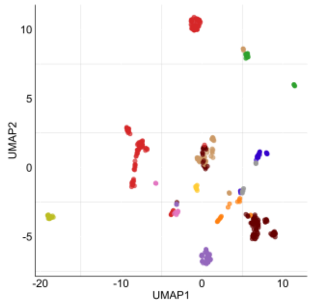 |
| MS1<br>presence/absence<br>rfe2 | 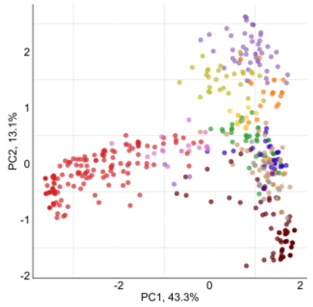 | 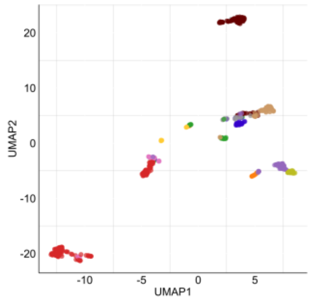 |
| MS1<br>log10<br>all | 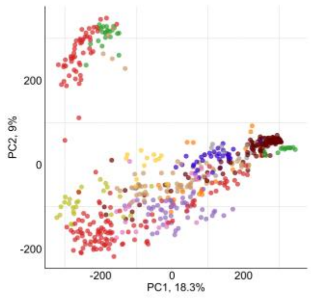 | 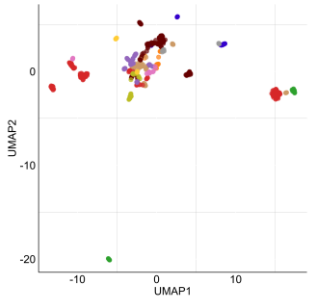 |

| | | |
|---|---|---|
| MS1<br>log10<br>rfe1 | 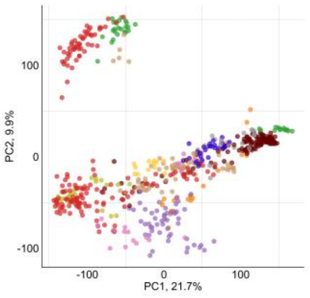 | 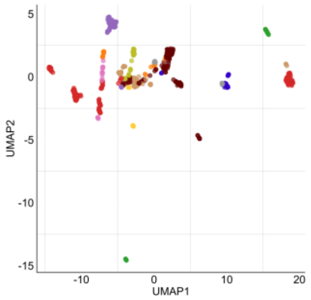 |
| MS1<br>log2<br>rfe2 | 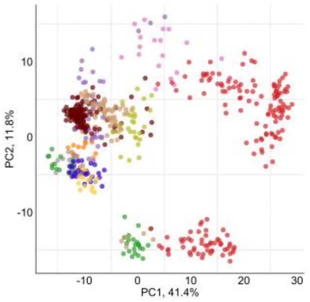 | 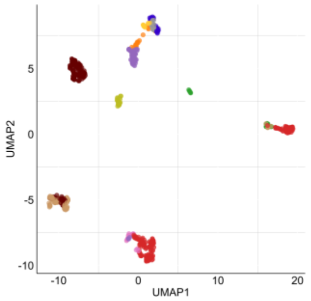 |

**Table S2.** PCA, UMAP for sample comparisons for MFP datasets. Principal component analysis (PCA), Uniform Manifold Approximation, and Projection (UMAP) for sample comparisons for molecular fingerprint (MFP) datasets. For chemical characteristic vectors, two different averaging was done. Regular averaged with all LC-MS features, and mass-grouped, where LC-MS features were separated in seven groups based on their MS1 precursor mass. For both approaches, datasets were analyzed with all features (marked as all), selecting all important features from recursive feature elimination 10-fold cross-validation (rfe1) and selecting consistently important features present in all 10-fold cross-validation when using recursive feature elimination (rfe2).

| | | |
|---|---|---|
| MFP<br>averaged<br>all | 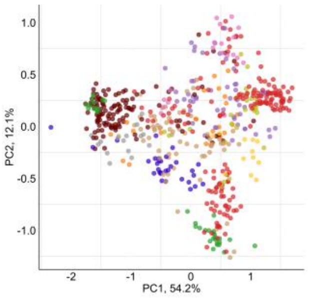 | 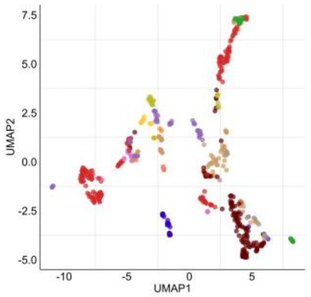 |
| MFP<br>averaged<br>rfe1 | 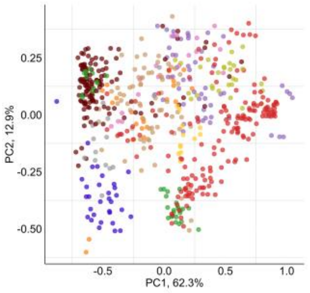 | 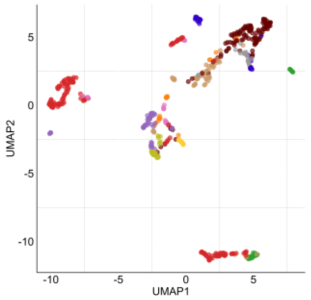 |

| | | |
|---|---|---|
| MFP<br>averaged<br>rfe2 | 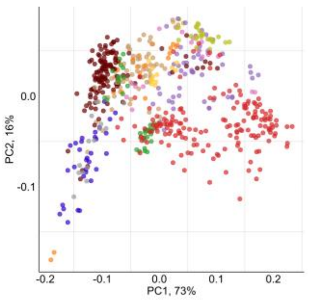 | 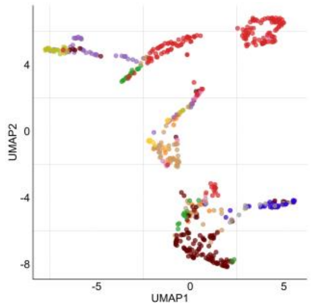 |
| MFP<br>mass grouped<br>all | 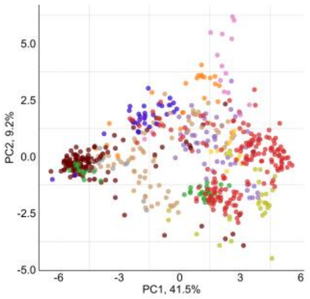 | 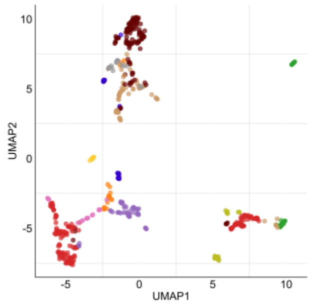 |
| MFP<br>mass grouped<br>rfe1 | 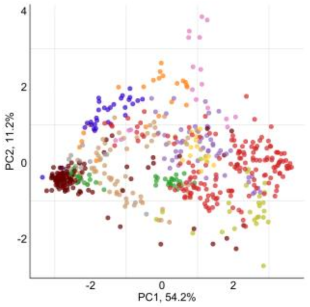 | 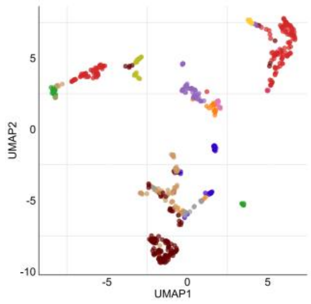 |
| MFP<br>mass grouped<br>rfe2 | 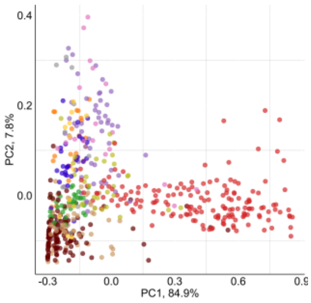 | 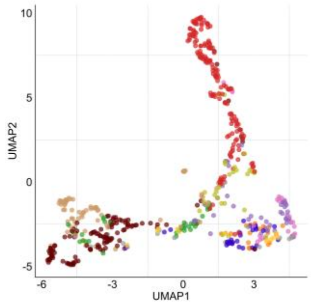 |

**Table S3.** Principal component analysis (PCA), Uniform Manifold Approximation, and Projection (UMAP) for sample comparisons for compound class (CC) datasets. For chemical characteristic vectors, two different averaging was done. Regular averaged with all LC-MS features, and mass-grouped, where LCMS features were separated into seven groups based on their MS1 precursor mass. For both approaches, datasets were analyzed with all features (marked as all), selecting all important features from recursive feature elimination 10-fold cross-validation (rfe1) and selecting consistently important features present in all 10-fold cross-validation when using recursive feature elimination (rfe2).

| | | |
|---|---|---|
| CC<br>averaged<br>all | 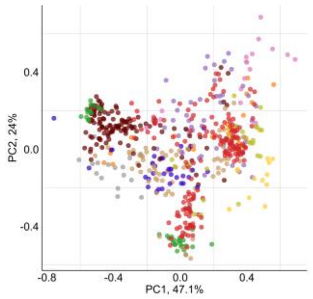 | 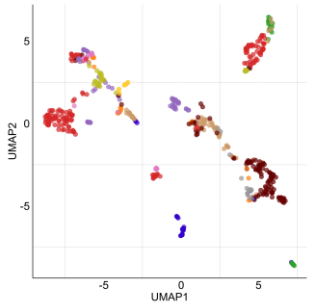 |
| CC<br>averaged<br>rfe1 | 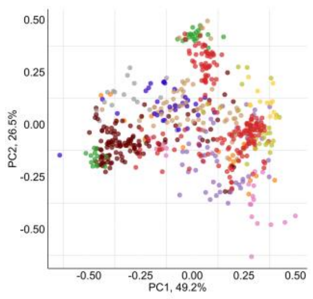 | 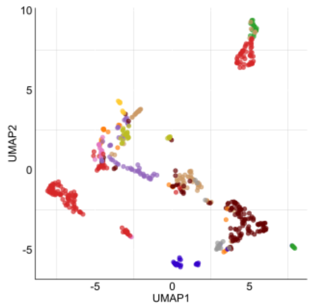 |
| CC<br>averaged<br>rfe2 | 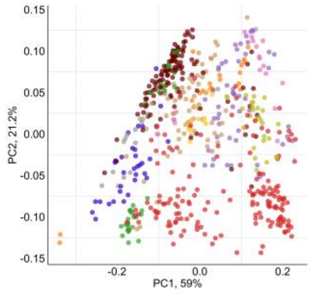 | 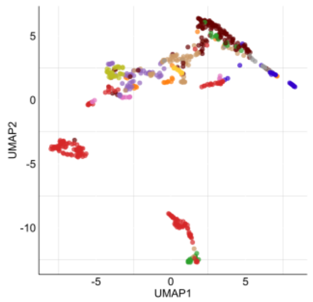 |
| CC<br>mass<br>all | 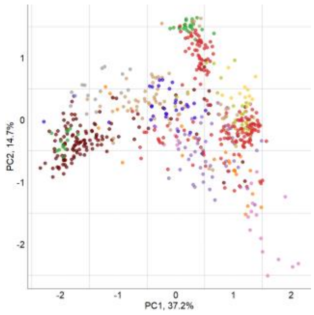 | 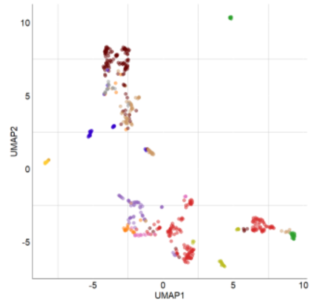 |

| | | |
|---|---|---|
| CC<br><br>mass<br><br>rfe1 | 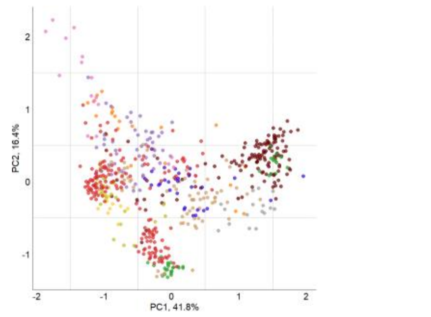 | 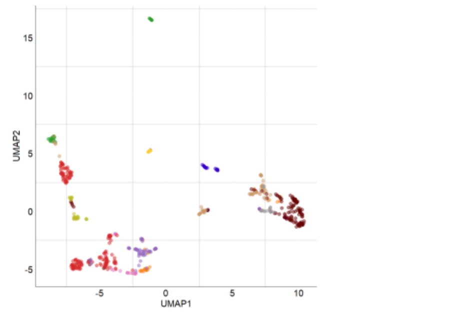 |
| CC<br><br>mass<br><br>rfe2 | 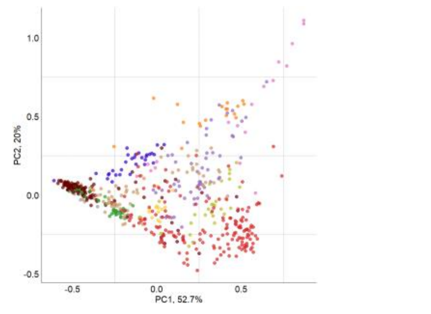 | 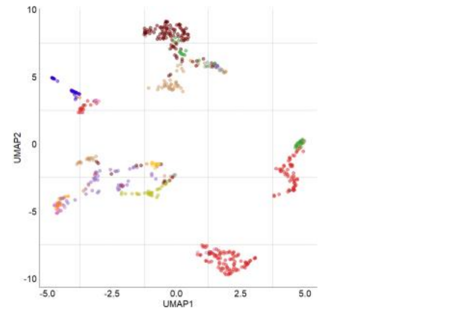 |

**Table S4.** The accuracy achieved for all methods with the k-Nearest Neighbors (kNN), Random Forest (RF) and Naive Bayes (NB) classifiers, as compared to the trivials, Random (Equal Probability) and Most Frequent. Methods are divided by the chemical characteristics (compound class - CC, molecular fingerprint - MFP), averaging approach with either averaging over all LC-MS peaks or using a mass-grouped approach with seven groups (marked as mass in the method name) and feature selection (all - all features selected excluding zero columns, rfe1 - all important features from recursive feature elimination 10-fold cross-validation, rfe2 - consistently important features present in all 10-fold cross-validation).

| Method | k-NN | RF | NB | Random | Most Freq. |
|---|---|---|---|---|---|
| CC_average_all | 0.90 | 0.86 | 0.95 | 0.09 | 0.29 |
| CC_average_rfe1 | 0.89 | 0.89 | 0.92 | 0.09 | 0.29 |
| CC_average_rfe2 | 0.88 | 0.89 | 0.86 | 0.09 | 0.29 |
| CC_mass_all | 0.92 | 0.84 | 0.96 | 0.09 | 0.29 |
| CC _mass_rfe1 | 0.92 | 0.90 | 0.93 | 0.09 | 0.29 |
| CC_mass_rfe2 | 0.89 | 0.90 | 0.82 | 0.09 | 0.29 |
| MFP_average_all | 0.91 | 0.89 | 0.93 | 0.09 | 0.29 |
| MFP_average_rfe1 | 0.92 | 0.91 | 0.92 | 0.09 | 0.29 |
| MFP_average_rfe2 | 0.82 | 0.86 | 0.78 | 0.09 | 0.29 |
| MFP_mass_rfe1 | 0.93 | 0.91 | 0.93 | 0.09 | 0.29 |
| MFP_mass_rfe2 | 0.82 | 0.81 | 0.68 | 0.09 | 0.29 |
| MFP_mass_all | 0.92 | 0.86 | 0.95 | 0.09 | 0.29 |
| MS1_log_all | 0.92 | 0.87 | 0.95 | 0.09 | 0.29 |
| MS1_log_rfe1 | 0.96 | 0.92 | 0.95 | 0.09 | 0.29 |
| MS1_log_rfe2 | 0.93 | 0.92 | 0.92 | 0.09 | 0.29 |
| MS1_presence_all | 0.96 | 0.78 | 0.94 | 0.09 | 0.29 |
| MS1_presence_rfe1 | 0.97 | 0.87 | 0.96 | 0.09 | 0.29 |
| MS1_presence_rfe2 | 0.94 | 0.89 | 0.87 | 0.09 | 0.29 |

55

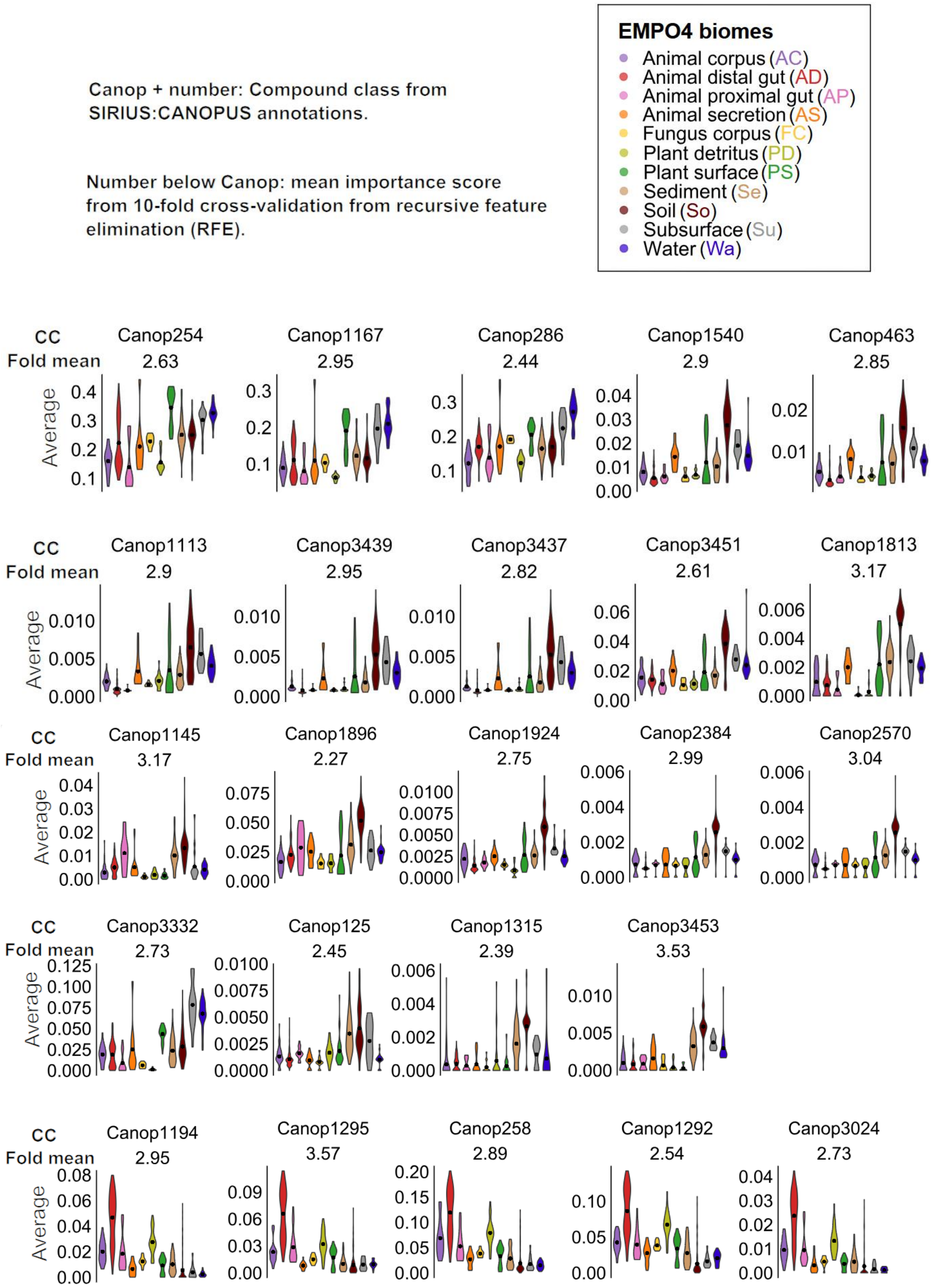

Canop + number: Compound class from SIRIUS:CANOPUS annotations.
Number below Canop: mean importance score from 10-fold cross-validation from recursive feature elimination (RFE).
EMPO4 biomes
Animal corpus (AC)
Animal distal gut (AD)
Animal proximal gut (AP)
Animal secretion (AS)
Fungus corpus (FC)
Plant detritus (PD)
Plant surface (PS)
Sediment (Se)
Soil (So)
Subsurface (Su)
Water (Wa)
CC
Fold mean
Average
Canop254 2.63
Canop1167 2.95
Canop286 2.44
Canop1540 2.9
Canop463 2.85
Canop1113 2.9
Canop3439 2.95
Canop3437 2.82
Canop3451 2.61
Canop1813 3.17
Canop1145 3.17
Canop1896 2.27
Canop1924 2.75
Canop2384 2.99
Canop2570 3.04
Canop3332 2.73
Canop125 2.45
Canop1315 2.39
Canop3453 3.53
Canop1194 2.95
Canop1295 3.57
Canop258 2.89
Canop1292 2.54
Canop3024 2.73

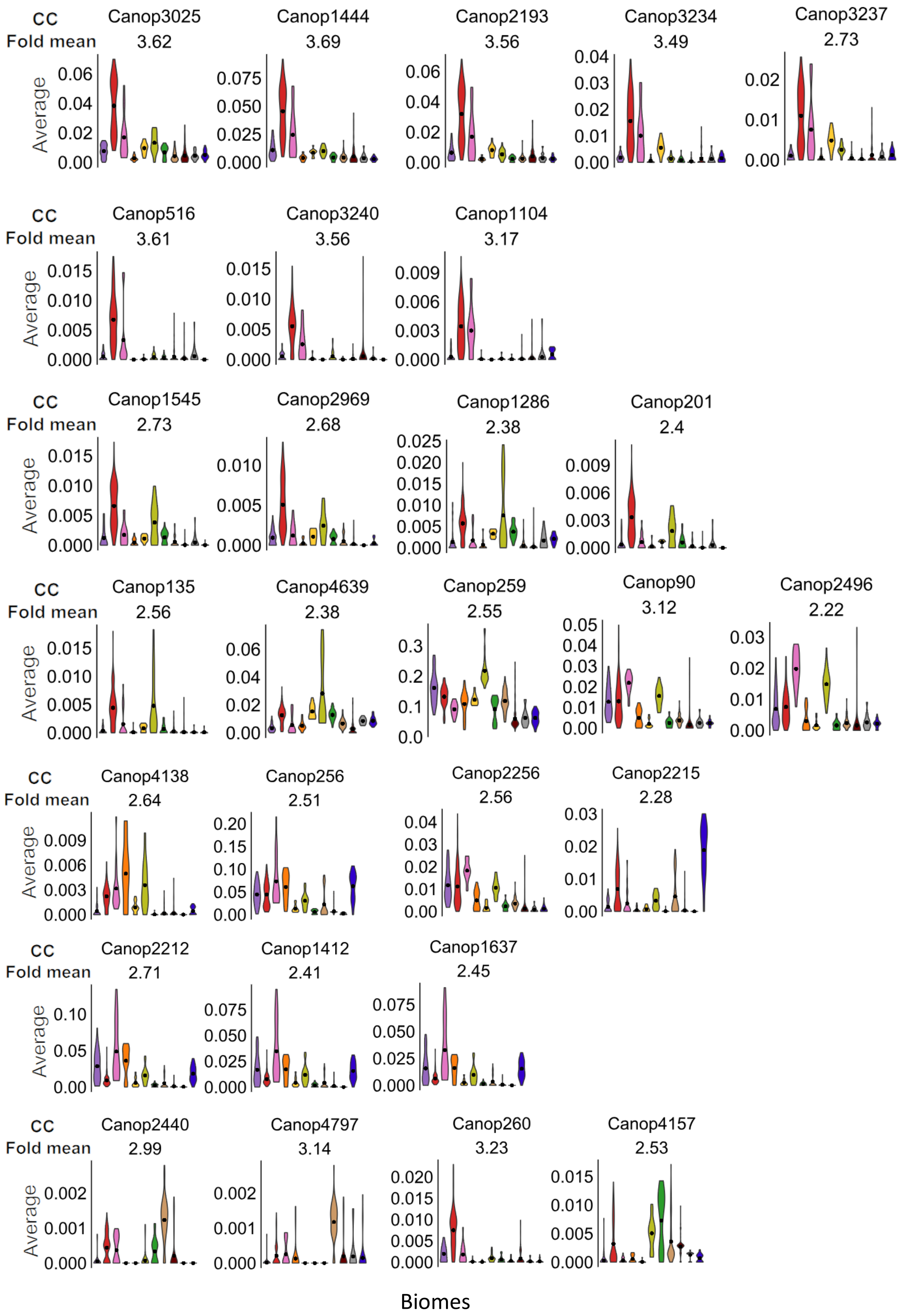


**Figure S4.** Selected 52 consistently important compound classes and their averaged values per sample grouped based on their assigned biomes. Compound class plots are manually organized based on their similarities. Each plot has its own scale since there are big differences between compound classes. Corresponding names for compound class codes from SIRIUS are presented in Table S5.

**Table S5.** Compound class codes from SIRIUS+CANOPUS and corresponding compound class names.

| Compound class | Compound class name |
|---|---|
| Canop254 | Ethers |
| Canop1167 | Dialkyl ethers |
| Canop286 | Primary alcohols |
| Canop1540 | Monosaccharide phosphates |
| Canop463 | Organic oxoanionic compounds |
| Canop1113 | Alpha-keto acids and derivatives |
| Canop3439 | Glycerone phosphates |
| Canop3437 | Glycerone and derivatives |
| Canop3451 | Monoalkyl phosphates |
| Canop1813 | Phenylphosphines and derivatives |
| Canop1145 | N-acylethanolamines |
| Canop1896 | 1,2-aminoalcohols |
| Canop1924 | Aminobenzenesulfonamides |
| Canop3332 | Polyethylene glycols |
| Canop125 | Quinomethanes |
| Canop1315 | 1,3-dioxolanes |
| Canop2384 | O-quinones |
| Canop2570 | Phenanthraquinones |
| Canop3453 | Trialkyl phosphates |
| Canop1194 | Oxosteroids |
| Canop1295 | Hydroxysteroids |
| Canop258 | Steroids and steroid derivatives |
| Canop1292 | Cyclic alcohols and derivatives |
| Canop3024 | 3-oxosteroids |
| Canop3025 | 3-hydroxysteroids |
| Canop1444 | Bile acids, alcohols and derivatives |
| Canop2193 | Hydroxy bile acids, alcohols and derivatives |
| Canop3234 | 7-hydroxysteroids |
| Canop3237 | 12-hydroxysteroids |
| Canop516 | Dihydroxy bile acids, alcohols and derivatives |
| Canop3240 | 7-oxosteroids |
| Canop1104 | Trihydroxy bile acids, alcohols and derivatives |
| Canop1545 | Vitamin E compounds |
| Canop2969 | 3-hydroxy delta-5-steroids |
| Canop1286 | Benzenediols |
| Canop201 | Tocopherols |
| Canop135 | Catechols |
| Canop4639 | 1-hydroxy-4-unsubstituted benzenoids |
| Canop4138 | L-alpha-amino acids |
| Canop256 | Glycerophospholipids |
| Canop2256 | Substituted pyrroles |
| Canop2215 | Glycerophosphoglycerols |

| | |
|---|---|
| Canop2212 | Glycerophosphocholines |
| Canop1412 | Lysophosphatidylcholines |
| Canop1637 | 1-acyl-sn-glycero-3-phosphocholines |
| Canop2440 | Naphthols and derivatives |
| Canop4797 | Furoic acid esters |
| Canop90 | Pyrroles |
| Canop2496 | Indoles |
| Canop259 | Prenol lipids |
| Canop260 | Monohydroxy bile acids, alcohols and derivatives |
| Canop4157 | Phenolic glycosides |

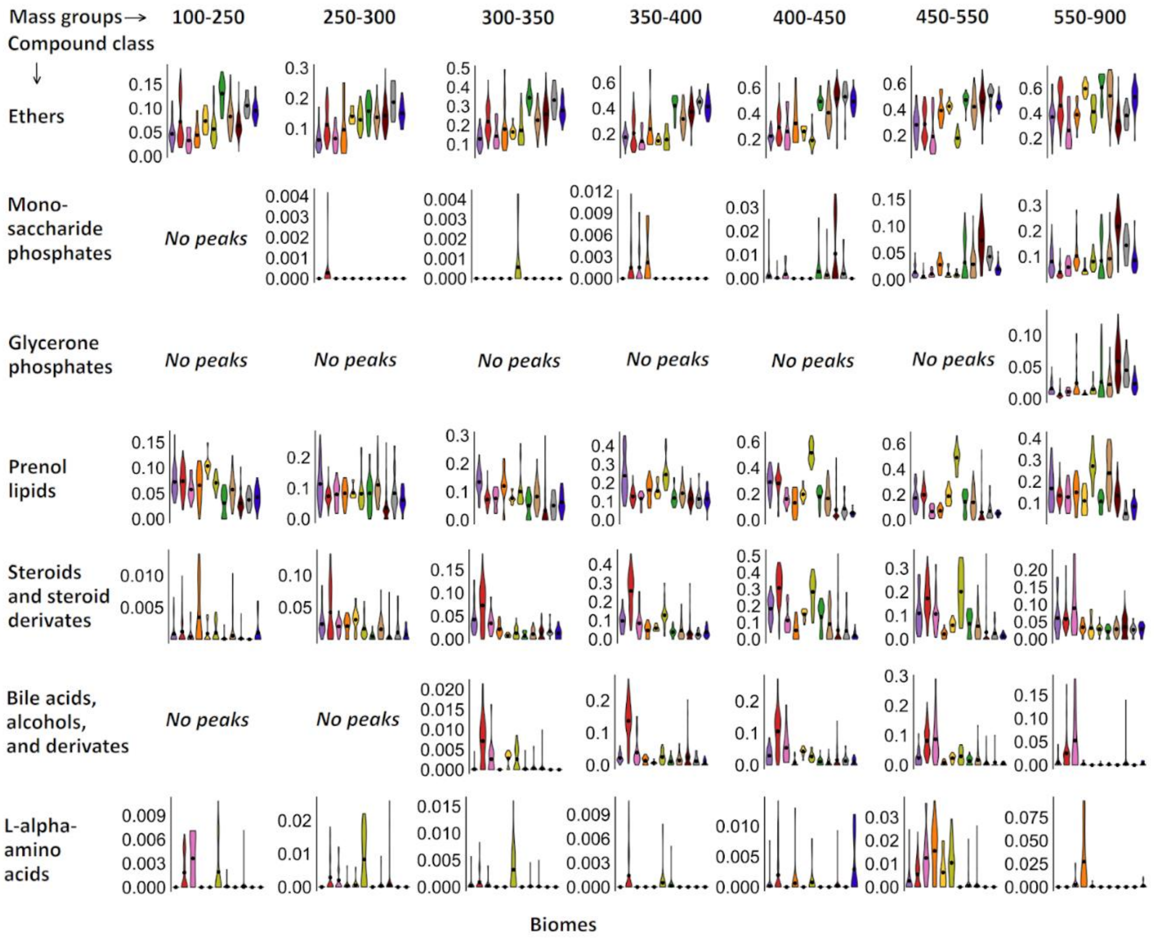


**Figure S5.** Seven selected chemical classes from the CCV approach, presented with corresponding averages from mass-grouped approach.

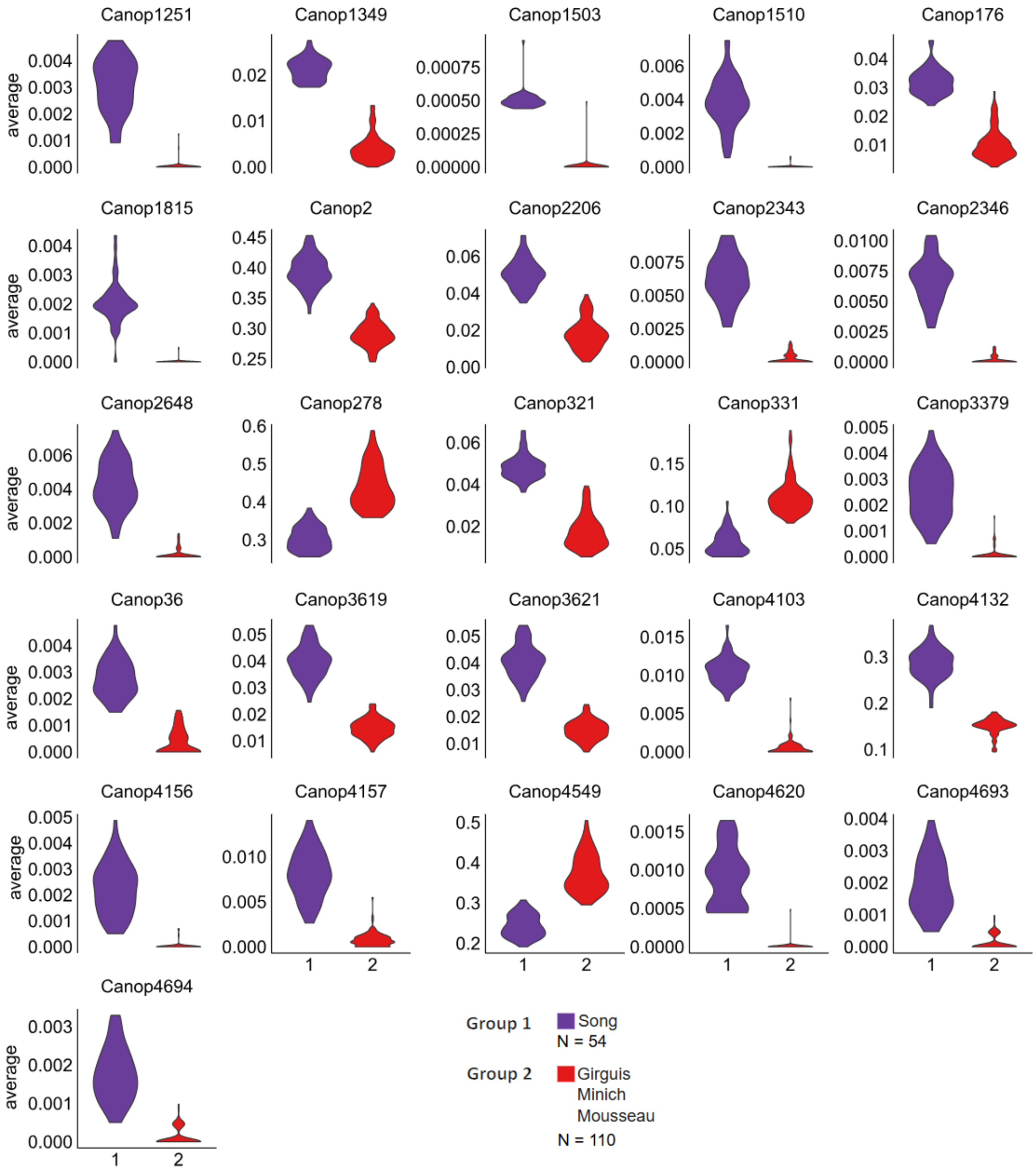


**Figure S6.** Animal distal gut samples were grouped into two based on UMAP clustering for within-biome analysis. Each plot shows on compound class and its ratio values for different samples. Compound class names for the CANOPUS (Canop) coding are in Table S6.

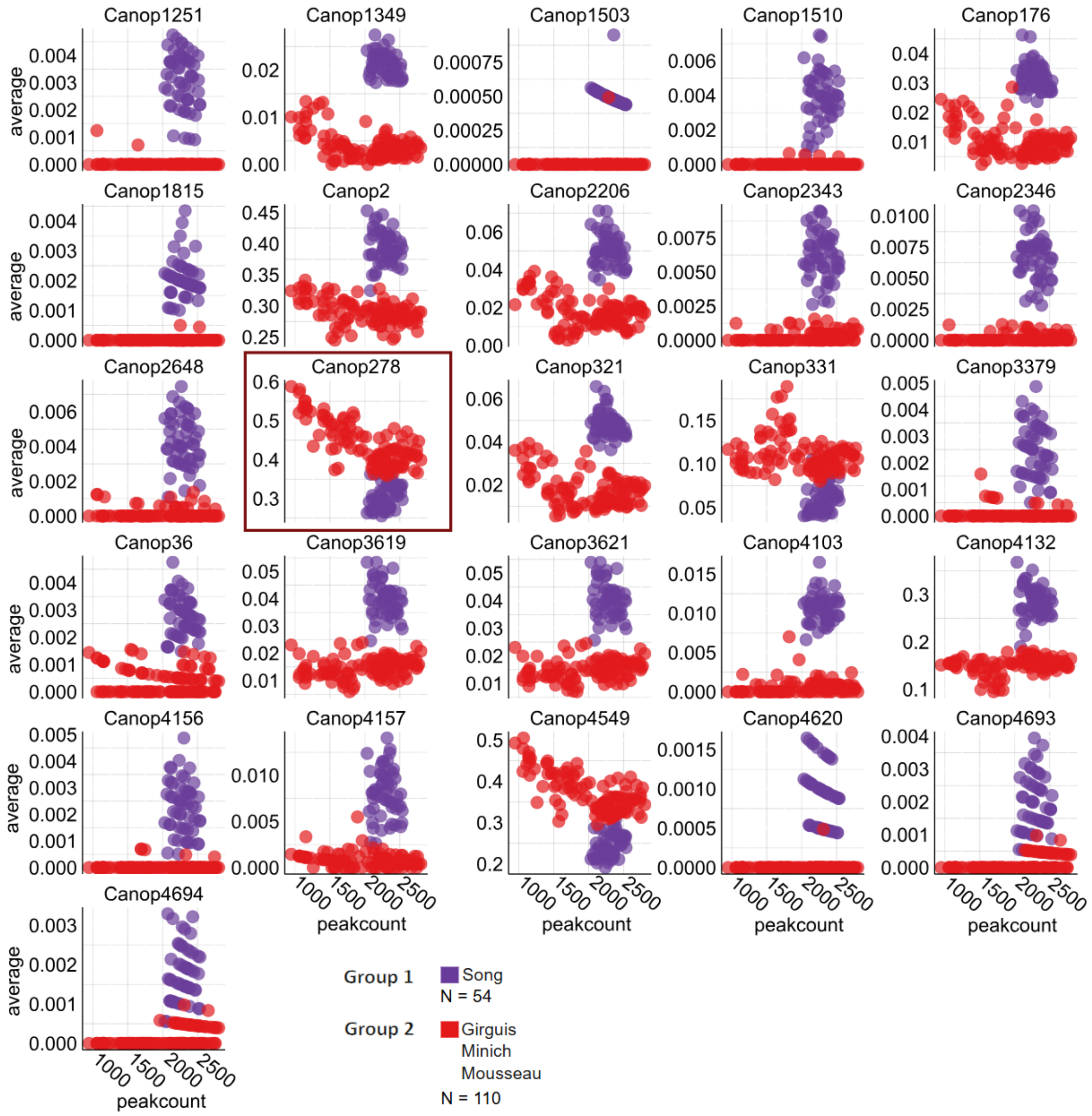


**Figure S7**. Correlation between averaged compound class values and amount of peaks that were used for the averaging. Each point on the graph is one sample from the animal distal gut biome. Canop278, Organonitrogen compounds plot, which is discussed more in the manuscript, is marked with a frame (third row, second column). Original data in Zenodo file "distalgut_groupanalysis.csv". Compound class names for the CANOPUS (Canop) coding are in Table S6.

**Table S6.** CANOPUS compounds class codes and names for Figure S6 and S7. Selected important compound classes to separate two animal distal gut groups.

| Compound class | Compound class name |
|---|---|
| Canop1251 | Gallic acid and derivatives |
| Canop1349 | Benzoic acid esters |
| Canop1503 | Diacylglycerophosphates |
| Canop1510 | Lignan glycosides |
| Canop176 | Benzoic acids and derivatives |
| Canop1815 | Furofurans |
| Canop2 | Organoheterocyclic compounds |
| Canop2206 | O-glycosyl compounds |
| Canop2343 | Methoxybenzoic acids and derivatives |
| Canop2346 | M-methoxybenzoic acids and derivatives |
| Canop2648 | Diarylheptanoids |
| Canop278 | Organonitrogen compounds |
| Canop321 | Benzoyl derivatives |
| Canop331 | Fatty amides |
| Canop3379 | Organic dithiophosphoric acids and derivatives |
| Canop36 | Nitrobenzenes |
| Canop3619 | Enoate esters |
| Canop3621 | Alpha,beta-unsaturated carboxylic esters |
| Canop4103 | Dimethoxybenzenes |
| Canop4132 | Oxacyclic compounds |
| Canop4156 | Dithiophosphate O-esters |
| Canop4157 | Phenolic glycosides |
| Canop4549 | Organopnictogen compounds |
| Canop4620 | 4-alkoxyphenols |
| Canop4693 | p-Hydroxybenzoic acid esters |
| Canop4694 | p-Hydroxybenzoic acid alkyl esters |

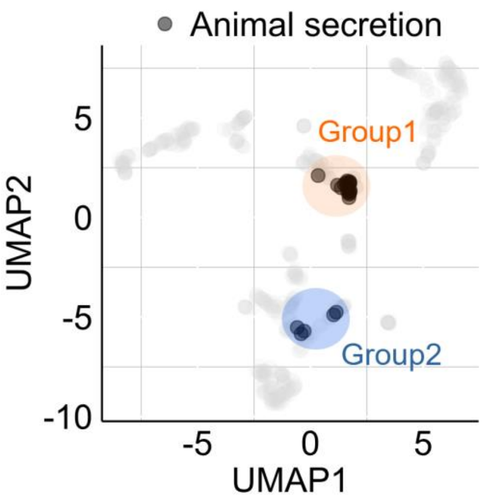


**Figure S8.** Animal secretion UMAP for mass-grouped molecular fingerprints (MFP). Groups for within-biome analysis were divided manually from this UMAP for the analysis of Figure 6e-h.

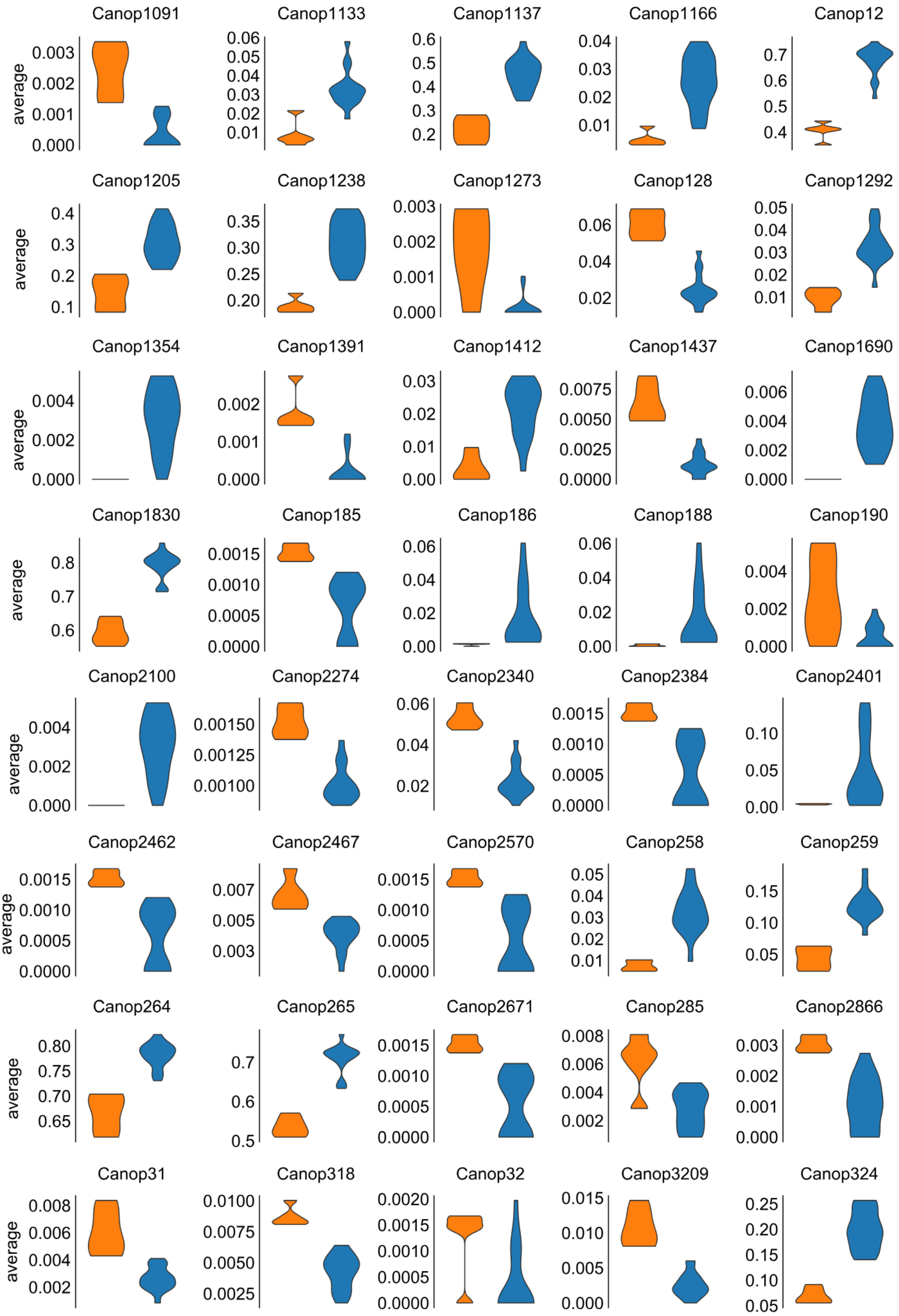
Canop1091
Canop1133
Canop1137
Canop1166
Canop12
Canop1205
Canop1238
Canop1273
Canop128
Canop1292
Canop1354
Canop1391
Canop1412
Canop1437
Canop1690
Canop1830
Canop185
Canop186
Canop188
Canop190
Canop2100
Canop2274
Canop2340
Canop2384
Canop2401
Canop2462
Canop2467
Canop2570
Canop258
Canop259
Canop264
Canop265
Canop2671
Canop285
Canop2866
Canop31
Canop318
Canop32
Canop3209
Canop324
average

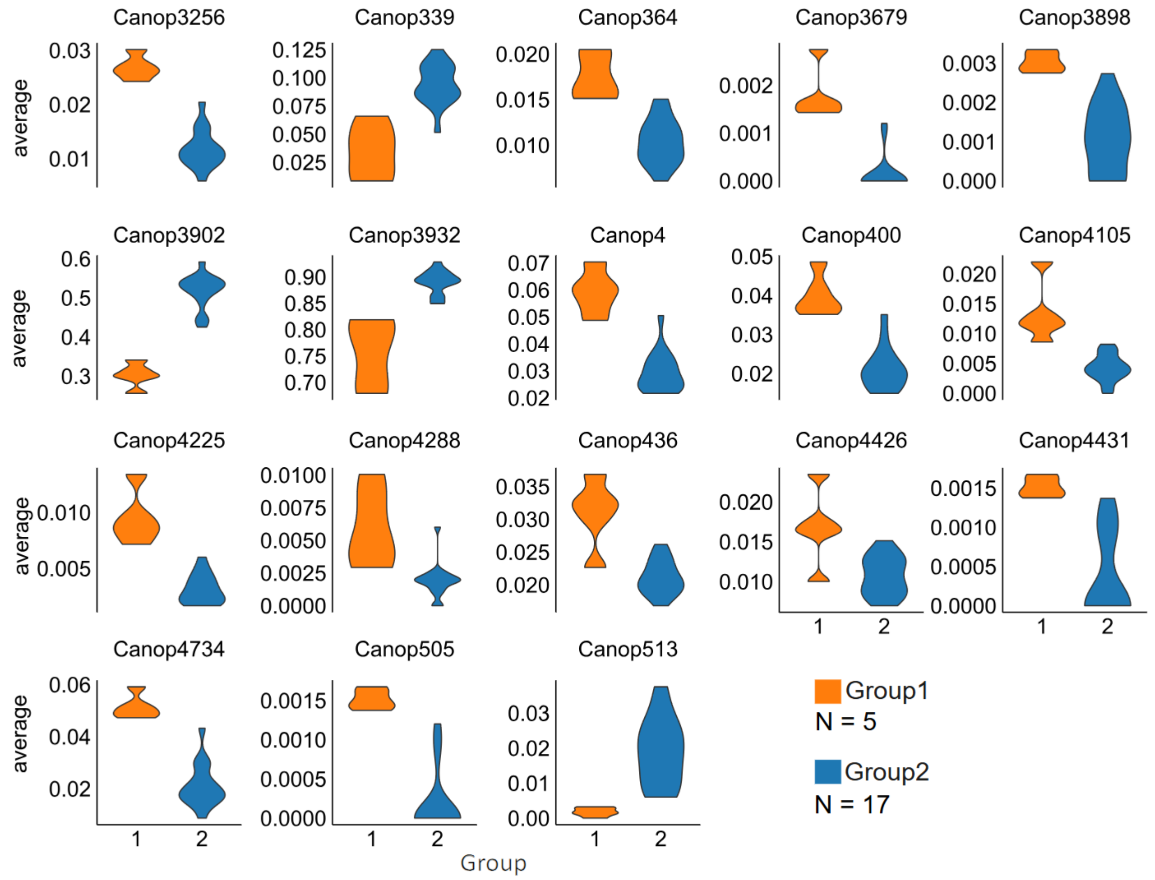


**Figure S9.** Coral samples (animal secretion) were grouped into two groups based on UMAP clustering for within-biome analysis (groups seen in Figure S7). Compound class names corresponding to CANOPUS coding (Canop) are in Table S7.

**Table S7.** CANOPUS compounds class codes and names for Figure S9. Selected important compound classes to separate two animal secretion biome groups.

| Compound class | Compound class name |
|---|---|
| Canop1091 | Toluenes |
| Canop1133 | Monoacylglycerols |
| Canop1137 | Monocarboxylic acids and derivatives |
| Canop1166 | Carboxylic acid salts |
| Canop12 | Lipids and lipid-like molecules |
| Canop1205 | Carboxylic acids |
| Canop1238 | Carboxylic acid esters |
| Canop1273 | Meta cresols |
| Canop128 | Alkyl aryl ethers |
| Canop1292 | Cyclic alcohols and derivatives |
| Canop1354 | Lysophosphatidylinositols |
| Canop1391 | Lignans, neolignans and related compounds |
| Canop1412 | Lysophosphatidylcholines |
| Canop1437 | Organothiophosphorus compounds |
| Canop1690 | Steroid esters |

| | |
|---|---|
| Canop1830 | Carbonyl compounds |
| Canop185 | Phenylmethylamines |
| Canop186 | Phenethylamines |
| Canop188 | Amphetamines and derivatives |
| Canop190 | Methoxyphenols |
| Canop2100 | 1-acyl-sn-glycerol-3-phosphoinositols |
| Canop2274 | Nitronaphthalenes |
| Canop2340 | Phenol ethers |
| Canop2384 | O-quinones |
| Canop2401 | N-acyl-alpha amino acids |
| Canop2462 | Diarylethers |
| Canop2467 | Organoselenium compounds |
| Canop2570 | Phenanthraquinones |
| Canop258 | Steroids and steroid derivatives |
| Canop259 | Prenol lipids |
| Canop264 | Organic acids and derivatives |
| Canop265 | Carboxylic acids and derivatives |
| Canop2671 | Benzylamines |
| Canop285 | Anilides |
| Canop2866 | Vinyl halides |
| Canop31 | Benzenesulfonamides |
| Canop318 | Carbamic acids and derivatives |
| Canop32 | Benzenesulfonic acids and derivatives |
| Canop3209 | Aryl-aldehydes |
| Canop324 | Fatty acid esters |
| Canop3256 | Sulfenyl compounds |
| Canop339 | Unsaturated fatty acids |
| Canop364 | Organic carbonic acids and derivatives |
| Canop3679 | Furanoid lignans |
| Canop3898 | Chloroalkenes |
| Canop3902 | Fatty Acyls |
| Canop3932 | Organic oxides |
| Canop4 | Organosulfur compounds |
| Canop400 | Organophosphorus compounds |
| Canop4105 | Methoxybenzenes |
| Canop4225 | Benzenesulfonyl compounds |
| Canop4288 | Phenylketones |
| Canop436 | Azoles |
| Canop4426 | Organic sulfonic acids and derivatives |
| Canop4431 | Organic sulfonamides |
| Canop4734 | Phenoxy compounds |
| Canop505 | Sulfones |
| Canop513 | Eicosanoids |
